# Unexpected excess heterozygosity in a threatened, mycoheterotrophic, self-pollinating orchid

**DOI:** 10.64898/2026.10.07.757398

**Authors:** Craig F. Barrett, Nicole M. Fama, John V. Freudenstein, Gerardo Salazar, Sandra J. Simon, Brandon T. Sinn

**Affiliations:** Department of Biology, West Virginia University, 53 Campus Drive, Morgantown, West Virginia, USA 26506; Wake Forest University School of Medicine, Bowman Gray Center for Medical Education, 475 Vine Street, Winston-Salem, North Carolina, USA 27101; Department of Evolution, Ecology, and Organismal Biology, Ohio State University, 1315 Kinnear Road, Columbus, Ohio, USA 43212; Instituto de Biología, Universidad Nacional Autónoma de México, Circuito Exterior, Ciudad Universitaria, Copilco, Coyoacán, A.P. 70-233 México, Distrito Federal. C.P. 04510; Biology and Biochemistry Department, Emory and Henry University, P.O. Box 947 Emory, Virginia, USA 24327; Department of Biology and Earth Science, Otterbein University, 1 South Grove Street, Westerville, Ohio, USA 43801; Faculty of Medicine and Life Sciences, University of Latvia, Jelgavas iela 3, Riga, LV-1004, Latvia

**Keywords:** *Corallorhiza*, Single nucleotide polymorphisms (SNPs), overdominance, extinction, selfing, mycoheterotrophy

## Abstract

Self-fertilization results in reduction of heterozygosity over time, accelerated by small population sizes, restricted ranges, and fragmented habitats. Yet some species maintain heterozygosity despite these factors. One is *Corallorhiza bentleyi*, a threatened, mycoheterotrophic, self-pollinating orchid restricted to Virginia and West Virginia, USA. Reduced representation sequencing was employed across five of the ∼12 known localities of *C*. *bentleyi*, including representatives of sister species *C. involuta* from Mexico, to quantify patterns of population structure and genomic diversity. Surveys of *C. bentleyi* revealed a predominance of the cleistogamous (closed-flowered) growth form. Differentiation was observed between the two species but with weak population structure in *C. bentleyi*. Surprisingly, excess heterozygosity and negative inbreeding were recovered in both species, counter to expectations. Analyses provided evidence against bioinformatic artifacts (collapsed paralogs), small-sample bias (Robertson effect), and accumulation of mutations along chromosomes due to low recombination associated with selfing (Meselson effect) as explanations for excess heterozygosity. Demographic analyses favored a model of post-glacial divergence among *C. bentleyi* and *C. involuta* lacking bottlenecks. Finally, simulations under alternative selection scenarios favored models of overdominance vs. those of negative frequency dependent selection. It was concluded that overdominance, namely selection against selfed homozygotes, is the most likely explanation for elevated heterozygosity in *C. bentleyi*. Conservation concerns are discussed in light of these findings and the threats of extinction for one of North America’s most vulnerable orchid species.

## 1. Introduction

Self-pollinating (selfing) plant species are expected to display reduced genetic diversity, particularly with respect to heterozygosity (East and Hayes, 1912; Wright, 1921; Stebbins, 1957; Charlesworth and Wright, 2003). Theory predicts that with each round of selfing, approximately half of heterozygotes are lost each generation (Mendel, 1866; Wright, 1921). Selfing populations eventually advance to near-complete homozygosity; an obligately-selfing, diploid population consisting of all heterozygotes is expected to reach >99.9% homozygosity in ten generations (Wright, 1921; Stebbins, 1957). Small population sizes and geographically isolated or fragmented habitats may amplify this effect (Templeton et al., 1990; Young et al., 1996). Erosion of heterozygosity is of great conservation concern because the lack of genetic diversity is often associated with low adaptive potential, inbreeding depression, and elevated extinction risk; namely, decreased fitness due to unmasked expression of deleterious recessive alleles (Barrett and Charlesworth, 1991; Charlesworth and Charlesworth, 1999; Frankham, 2005).

While the situation above may be the case for many species, others present a conundrum: how do some species maintain excess heterozygosity, contrary to theoretical expectations, despite clonality, inbreeding, or selfing? Notable examples include *Prunus avium* (L.) L., *Populus tremuloides* Michx., and *Taraxacum officinale* F.H. Wigg. (Stoeckel et al., 2006; McAssey et al., 2017; Hernández-Velasco et al., 2024). Mechanisms like polyploidy, in which additional genome copies and subsequent mutation confer increased genomic diversity, necessitate additional generations to reach total homozygosity, and therefore provide a means to escape deleterious heterozygosity loss (Stebbins, 1950; Moody et al., 1993; Soltis and Soltis, 2000; Comai, 2005; Doyle et al., 2008; Parisod et al., 2010). However, even without polyploidy, there may be selection against homozygotes (overdominance) or negative frequency dependent selection (NFDS) in inbred or selfing populations if rare alleles confer fitness advantages over space-time, both of which may manifest as balancing selection (e.g., Fisher, 1930; Wright, 1931; Takahata and Nei, 1990; Hedrick, 2012; Allendorf et al., 2017; Hartfield et al., 2017). Further, the Meselson Effect, a phenomenon that involves selfing populations experiencing mutations that accumulate among non-recombining chromosome copies, may lead to apparent increases in heterozygosity as the chromosome copies diverge, as well as extensive haplotype structure via linkage (Welch and Meselson, 2000).

Bentley’s coralroot (*Corallorhiza bentleyi* Freudenst.) is a rare mycoheterotrophic orchid known from a dozen sites in Appalachia, and an apt species in which to study the consequences of selfing and rarity. It was discovered in 1996 and subsequently described as a new species of *Corallorhiza* (Freudenstein, 1999; Bentley, 2004). A census in 2001 estimated a total population size of ∼400 individuals among known localities in Virginia and West Virginia, USA (Bentley, 2004). The International Union for Conservation of Nature (IUCN) Red List classifies *C. bentleyi* as “D1-Vulnerable,” and NatureServe lists *C. bentleyi* as “G2-Imperiled” (Goedeke et al., 2015; NatureServe, 2026). Subsequent late-summer surveys of the largest historically known population in Giles County, Virginia, USA from 2016-2024 yielded no racemes, underscoring the threat of extinction to this species (Barrett, C. F., *personal observation*).

*Corallorhiza bentleyi* is a non-photosynthetic parasite that specializes upon a narrow suite of ectomycorrhizal fungal hosts in the family Thelephoraceae Chevall. for all nutritional needs, representing a major biotic niche restriction in this orchid to specific forest habitats (Barrett et al., 2010; Fama et al., 2021). This species displays characteristic evidence of obligate self-pollination (selfing syndrome; Darwin, 1876; Ornduff, 1969; Richards, 1986; Sicard and Lenhard, 2011). Its drably colored flowers are highly reduced, with all flowers having swollen ovaries and seed-set at anthesis, though the viability of selfed seeds is unknown. Plants from the type locality in Monroe County, West Virginia, USA were originally described as cleistogamous (closed-flowered; Fig. 1), though subsequent observations of this species revealed that some populations contain individuals that are slightly more open-flowered (Bentley, 2004; Fig. 1). Our observations in the field suggest that ovary swelling and seed development begin *prior* to anthesis, even in more open-flowered forms. The restricted geographic range, disjunction from its sister species, specialized fungal associations, low census population size estimates, and predominant or obligate mode of self-pollination make *Corallorhiza bentleyi* an excellent target system in which to test predictions for the downstream effects of these traits on genetic diversity.

**Figure 1.**
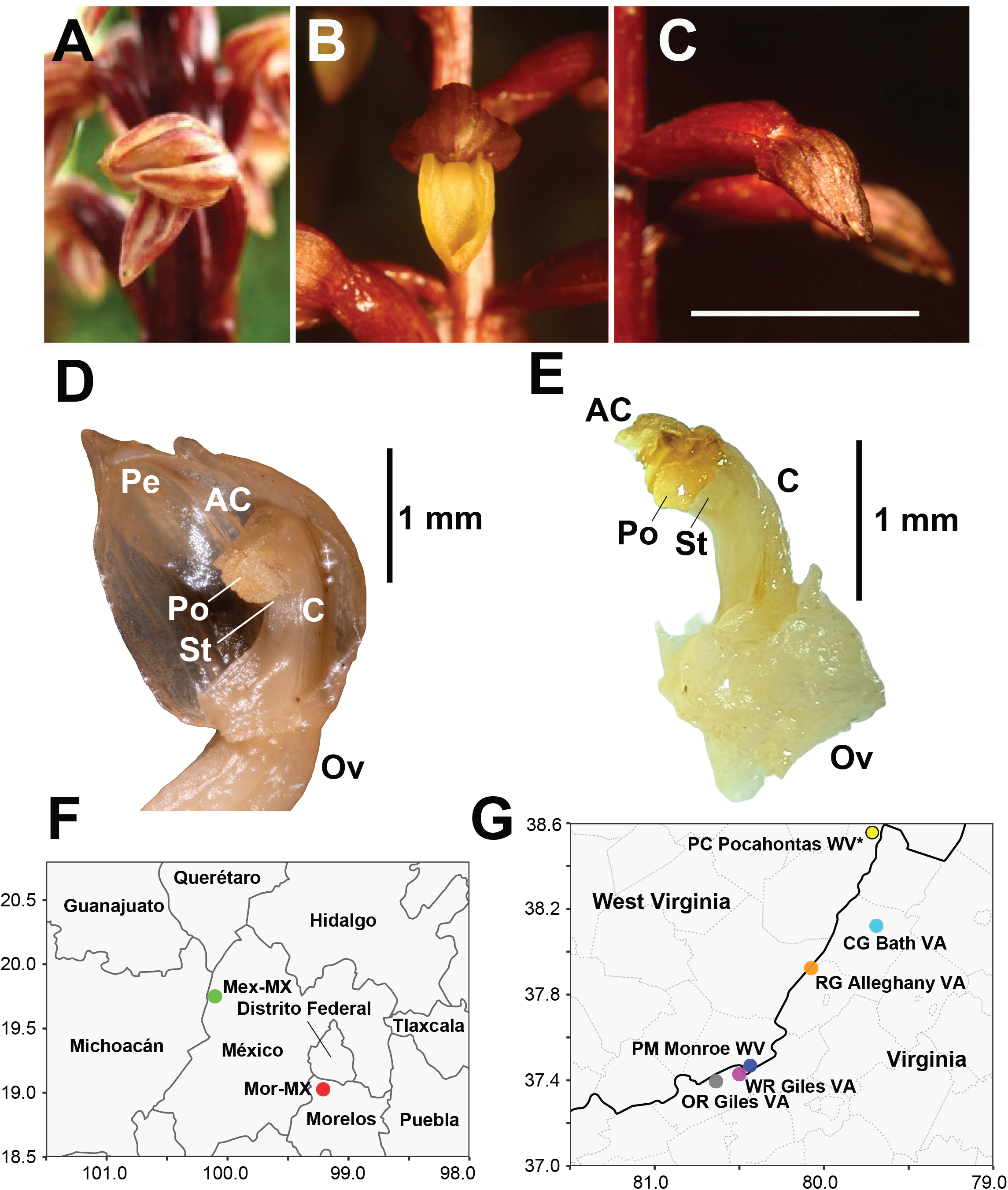
**A**. *Corallorhiza involuta*, Morelos, Mexico. **B**. Chasmogamous flower of *C. bentleyi*, Monroe County, West Virginia, USA. **C**. Cleistogamous flower of *C. bentleyi*, Monroe County, West Virginia, USA. Scale bar = 1 cm. **D**. Liquid-preserved flower of *C. involuta* (Oaxaca, Mexico) and **E**. *C. bentleyi* (Bath County, Virginia, USA), showing anther cap (AC), swollen pollinium (Po) attached to the stigma (St), distal end of the ovary (Ov), column (C), and parts of the perianth (Pe, *C. involuta* only). Scale bars = 1 mm. **F**. Map of two sampling localities of *C. involuta* in Mexico. Red circle = population 228 Morelos (Mor-MX), green circle = population 237 Mexico (Mex-MX). **G**. Map of two sampling localities of *C. bentleyi* in the eastern USA. ‘*’ Pocahontas County, West Virginia site (PC, yellow point) was not sampled for genomic data. Photos: C. Barrett, J. Freudentein.

*Corallorhiza bentleyi* is a member of the *Corallorhiza striata* Lindl. species complex, encompassing two varieties: the large-flowered, predominantly outcrossing *C. striata* var. *striata* Lindl. (western and northern US), and the smaller-flowered, predominantly selfing *C. striata* var. *vreelandii* Rydb. (southwestern US and Mexico; Freudenstein, 1997; 1999). *Corallorhiza involuta* Greenm., was previously treated as a variety of *C. striata* (var. *involuta*) but is now recognized as a distinct species and is somewhat broadly distributed in Mexico, though no formal conservation assessment exists for *C. involuta* (Freudenstein, 1997; Barrett and Freudenstein, 2011). This species most closely resembles *C. bentleyi* based on miniscule flower size, (presumably) predominant selfing, and a similar late-summer phenology (Freudenstein, 1999; Fig. 1). The main feature differentiating *C. bentleyi* from *C. involuta* is the extension of the callus apex (fused lamellae at the lip base) beyond the widest point of the lip, whereas this is not the case for all other members of the complex (Barrett and Freudenstein, 2011). Molecular data demonstrated a sister relationship between *C. involuta* and *C. bentleyi*, both of which collectively are divergent from the rest of the complex, but previous studies were based on one-to-few individuals of each species (Barrett and Freudenstein, 2009; 2011; Barrett et al., 2014; 2025). It was previously hypothesized that the restricted range of *C. bentleyi* represents a relict of a once-widespread ancestor of it and *C. involuta*, possibly in the southeastern US and northern Mexico during the Last Glacial Maximum (Barrett and Freudenstein, 2009; 2011).

A previous study based on dominant ISSR (Inter-Simple Sequence Repeat) banding patterns in *C. bentleyi* suggested a higher proportion of genetic diversity within populations than between them, the opposite of what was expected (Fama et al., 2021). The authors discussed the possibility that the patterns could have been the result of frequent seed dispersal among sites, refugial polymorphism following glacial retreat, population admixture following recent colonization of maturing secondary forest, homoplasy in ISSR banding patterns, or genotype-level symbiotic specificity upon fungal host lineages. However, the use of ISSR limited what conclusions could be drawn to explain these patterns, as they did not allow quantification of heterozygosity.

Here, we sampled from across the range of *C. bentleyi* including representatives of the Mexican *C. involuta*, employing genome-scale data from reduced representation sequencing (ISSR-seq; Sinn et al., 2021) with the following objectives: 1) determine relationships between these two taxa, and evidence for their distinctness; 2) characterize overall patterns of population structure between *C. involuta* and *C. bentleyi*, and especially within *C. bentleyi*; 3) quantify relative proportions of cleistogamous (closed-flowered) forms across localities, among individuals, and within individuals of *C. bentleyi*; 4) evaluate evidence for reduced genetic variation, specifically heterozygosity; and 5) determine the most plausible mechanism of selection with respect to heterozygosity. We hypothesize that *C. bentleyi* and *C. involuta* are distinct taxa, that population structure will be pronounced between the two species and within *C. bentleyi* due to its habit of obligate selfing, and that heterozygosity will be low in *C. bentleyi*, displaying a hallmark genomic signature of rarity and selfing.

## 2. Materials And Methods

### 2.1 Field assessment of chasmogamy-cleistogamy in *C. bentleyi*

Six *C. bentleyi* sites were surveyed over four years (2022-2025). These included two in Giles County, Virginia, USA (OR and WR), one in Monroe County, West Virginia, USA (PM), two in Bath County, Virginia, USA (CG and DK), and one in Pocahontas County, West Virginia, USA (PC; see map in Fig. 1). Not all sites were visited each year due to time and access constraints.

At each site, the total number of racemes was recorded, as were the numbers of cleistogamous and chasmogamous flowers per individual raceme. The objective was to determine whether plants at each locality were exclusively chasmogamous, cleistogamous, or mixed, and whether frequencies of chasmogamy-cleistogamy were consistent across years. To search for evidence of cleistogamy in *C. involuta*, herbarium visits and digitized specimen examinations were made. Visits were made in 2023 to the Oakes Ames Orchid Herbarium (AMES, *N* = 11 specimens) at Harvard University and the Smithsonian National Museum of Natural History Herbarium (US, *N* = 2 specimens). Searches for digitized specimens were conducted for US herbarium consortia (https://swbiodiversity.org, accessed 08 August, 2026). A search was also conducted in iNaturalist for all *Corallorhiza* in Mexico (https://www.inaturalist.org). To more closely explore the mechanism of self-pollination, we examined liquid-preserved material for representative specimens of both species.

### 2.2. Tissue sampling, DNA extraction, and ISSR sequencing

A single flower-ovary pair was carefully extracted from the multi-flowered raceme of each individual plant, so as not to destroy whole plants, with a minimum distance of 5m between individual plants. A 0.1g sample of ovary tissue was removed with a 100% ethanol-sterilized scalpel for DNA extraction using a 1/10-volume CTAB protocol (Doyle and Doyle, 1987). The perianth was removed and fixed in formalin (10% volume)-acetic acid (5% volume)-ethanol (50% volume) and transferred to 70% ethanol after three days for long-term storage. Whole-plant vouchers were not collected due to the rarity of both *C. involuta* and *C. bentleyi*; instead, spirit-preserved flowers serve as vouchers at the West Virginia University Herbarium (WVA). DNA quantification, ISSR amplification, and library preparation were carried out as in Sinn et al. (2021). ISSR amplicons were sequenced on an Illumina MiSeq (V2 chemistry, 2 × 150 bp paired-end reads) at the WVU Genomics Core Facility.

### 2.3. Genomic data assembly and filtering

ISSR loci were assembled following a dedicated ISSR assembly pipeline composed of three BASH scripts (ISSRseq_AssembleReference.sh, ISSRseq_CreateBams.sh, and ISSRseq_AnalyzeBAMs.sh; https://github.com/btsinn/ISSRseq/wiki). Briefly, the pipeline: trims reads based on quality and removes adapter sequences; builds a pseudoreference assembly from the accession with the most reads (here B8, Bath County, Virginia, USA); hard-trims flanking ISSR motifs from the resulting contigs; conducts BLAST searches of pseudoreference contigs against a database of plant, animal, and microbial genomes and removes contaminant contigs; maps reads to the pseudoreference; sorts the resulting BAM files; removes duplicate reads; and calls biallelic SNP variants (Sinn et al., 2021; Barrett et al., 2022).

We generated five primary datasets based on species inclusion (*C. bentleyi* only vs. *C. bentleyi* and *C. involuta*), missing-data filtering, and linkage thinning, and converted each to a genind object in R with adegenet v.2.1.11 (Jombart et al., 2008) (Table 1). First, the dataset comprised 52,718 SNPs (“gid_bent_unpruned”) for *C. bentleyi* after initial removal of the *C. involuta* samples. To prevent potentially duplicated loci from biasing diversity statistics, we removed 2,942 putative paralogs exhibiting excessive individual heterozygosity (> 85%), yielding a dataset of 49,776 SNPs (“gid_bent_cleaned”). This dataset was used to evaluate locus-level heterozygosity metrics, inbreeding coefficients, and individual multilocus heterozygosity. Second, we filtered the initial joint dataset containing both *C. bentleyi* and *C. involuta* for missing data (< 20%) in TASSEL v5, resulting in 15,958 variants (“gid”). To satisfy the assumption of marker independence for phylogenetic and population structure analyses, we pruned linked sites in VCFtools using a physical distance of one SNP per 1,000-bp window (--thin 1000). This produced a dataset of 2,347 SNPs (“gid_full”), which was analyzed both with and without the *C. involuta* accessions.

**Table 1.**
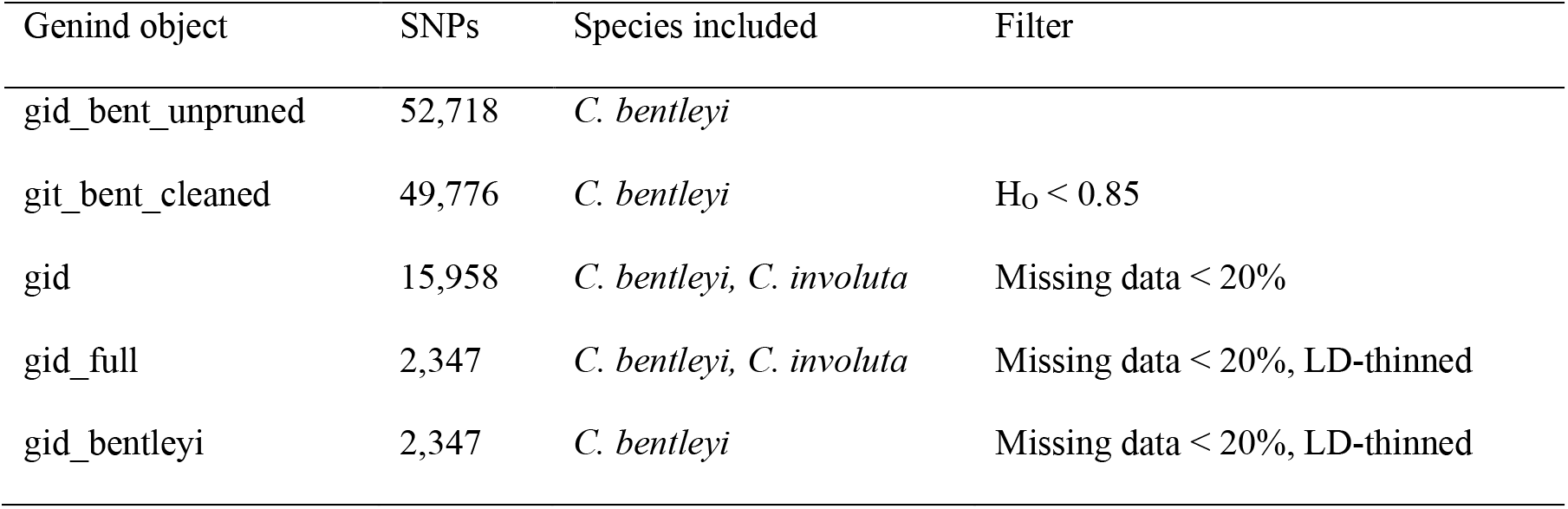
Summary of genind object datasets used for genomic data analyses. ‘SNPs’ = the number of biallelic single nucleotide polymorphisms. ‘Filter’ = the filtering method and parameters used for each dataset.

### 2.4. Relationships and population structure among *C. bentleyi* and *C. involuta*

The LD-thinned data (gid_full) were converted to NEXUS format using vcf2phylip v.2.8 (Ortiz, 2019; https://doi.org/10.5281/zenodo.2540861) and further processed with the R package phrynomics v.2.0, removing invariant, non-binary, and uninformative sites, as is required for downstream phylogenetic analyses to avoid ascertainment bias (Leaché et al., 2015). Nucleotide diversity (π) was calculated both within and between *C. involuta* (N = 5) and *C. bentleyi* (N = 37). Because of the sampling imbalance between the two species, we computed π ten times by randomly subsampling five individuals of *C. bentleyi* per replicate and used the mean and standard deviation for within-*C. bentleyi* estimates. A phylogenetic tree was computed in IQTREE3 v.3.1.3 (Nguyen et al., 2015; Wong et al., 2025) choosing the best-fit model for the concatenated SNP dataset and applying the ascertainment bias model (-m MFP+ASC). 1,000 ultra-fast bootstrap pseudoreplicates were conducted to assess branch support (Minh, 2013). The results were plotted with ggtree v.3.10.1 in R (Xu et al., 2012). Principal components analysis (PCA) on the gid_full and gid_bentleyi data was conducted in TASSEL5 based on the variance-covariance matrix, keeping the top five PCs, and plotted with ggplot2 v.4.0.3 in R (Wickham et al., 2016).

To investigate the degree of population structure and ancestry, sparse Non-negative Matrix Factorization (sNMF) on the LD-thinned gid_bentleyi dataset was conducted in the R package LEA v.3.19.8 (Frichot et al., 2014; Frichot and François, 2015) for *K* = 1-6, where *K* is the number of ancestral genomic clusters, using default alpha and tolerance values. The optimal *K*-value was determined via cross-entropy (Frichot et al., 2014). Discriminant Analysis of Principal Components was conducted in the adegenet for the gid_bentleyi data for *K* = 1-6, using both unsupervised (no population prior; “find.clusters”) and supervised analyses (with sampling localities serving as population priors). In the unsupervised analysis, the number of PCs retained was based on the plateau value of total variance explained, and the optimal K-value was chosen via the Bayesian Information Criterion. The A-score criterion was used to determine the optimal number of PCs in the supervised analysis (Jombart et al., 2010). These non-model-based methods were chosen over alternative methods such as STRUCTURE (Pritchard et al., 2000) and ADMIXTURE (Alexander et al., 2009) because they require no assumptions associated with Hardy-Weinberg Equilibrium or clonality.

To further characterize fine-scale population structure and quantify shared genomic ancestry without relying on allele frequency assumptions, we used fineRADstructure v.1.0 (Malinsky et al., 2018). Shared haplotype matrices were inferred in RADpainter v.1.0 (Malinsky et al., 2018) for the *C. bentleyi-*only dataset (gid_bentleyi). Hierarchical clustering and population structure were then inferred with fineSTRUCTURE v.4 (Lawson et al., 2012) using 100,000 MCMC steps, and a 100,000-step burn-in. A principal coordinates analysis (PCoA) on the co-ancestry matrix was conducted in R, with distances converted to a dissimilarity matrix [*D =* max(*coancestry*) *-coancestry*]. Analysis of Molecular Variance (AMOVA; Excoffier et al., 1992) was conducted on the gid_full dataset with the poppr v.2.9.8 R package (Kamvar et al., 2014), and pairwise Weir and Cockerham’s (1984) *F_ST_* analysis was conducted with the R package hierfst v.0.5-11 (Goudet, 2005).

### 2.5. Analyses of genomic diversity, inbreeding, and linkage

Basic population-genetic statistics were calculated with poppr and hierfstat globally and for each locality on the gid_full dataset: observed heterozygosity (*H_O_*), expected heterozygosity (*H_E_*), the ratio of *H_O_/H_E_*, and the inbreeding coefficient, *F_IS_*. Isolation-by-distance was then investigated via a Mantel test between genetic distance, calculated with adegenet, and geographic distance in R with 999 permutations. Poppr was used to convert gid_full into a genclone R-object and quantify the degree of linkage disequilibrium globally and within each population with the Index of Association (*I_A_*) and Standardized Index of Association metrics (*r̄_D_*, standardized by the number of loci), with 199 randomizations. Private alleles per species and per sampling locality were calculated in poppr with the ‘private_alleles’ function.

### 2.6. Data quality, filtering, and bioinformatic artifact investigations

To evaluate whether bioinformatic artifacts are associated with *Ho*, we examined the distribution of locus-level *Ho* values across 52,718 loci calculated using hierfstat from the gid_bent_unpruned dataset. Collapsed paralogs are expected to generate a prominent peak near *Ho* = 1 (i.e., apparent fixed heterozygosity). To screen putative paralogs, we calculated the proportion of heterozygous individuals at each locus. Loci showing near-fixed heterozygosity (greater than or equal to 85% of sampled individuals being heterozygous) were flagged as potential paralogs and removed, resulting in the gid_bent_cleaned dataset. Excess heterozygosity was quantified as Δ*H* = *Ho* - *H_E_* for each locus before and after putative paralog removal. A one-sample t-test was conducted on Δ*H* to determine whether a positive shift in heterozygosity persisted in putatively single-copy loci.

We tested the relationship between minor allele frequency (*MAF*) and *F_IS_* across all loci using hierfstat for both the gid_bent_unpruned and gid_bent_cleaned datasets with linear regression. Under selection against homozygotes, common variant sites (high *MAF* sites) are expected to experience selection against homozygous offspring, resulting in a negative slope (*β*), whereas collapsed paralog artifacts are expected to yield a flat relationship across *MAF* values (Hedrick, 2012). A Student’s t-test on the slope parameter *β*_observed_ was performed against the null expectation (*β* = 0) for both LD-pruned and unpruned data.

We investigated whether assembly methods influenced heterozygosity and inbreeding calculations. As an alternative to the pseudoreference-based approach (ISSRseq_AssembleReference.sh), we used discoSNP, a reference-free assembly pipeline that generates a VCF file directly from raw read data (Uricaru et al., 2015). We first concatenated the trimmed forward and reverse reads into single datasets for each accession, then ran the pipeline (run_discoSnp++.sh; kmer size = 31, minimum coverage depth = 10, maximum depth = 2,000). The resulting vcf file was further filtered in TASSEL5 to remove sites with > 20% missing data, and the same basic metrics as above were calculated in R (e.g., *Ho*, *H_E_*, *Ho*/*H_E_*, *F_IS_*) and compared to the original values from the pseudoreference assembly.

### 2.7. Empirical tests for heterozygosity excess and inbreeding

To test for sampling artifacts in small populations, where finite sampling may be expected to result in negative *Fis* values, we evaluated population-level *Fis* against small-*N* expectations for the gid_bent_cleaned dataset (E[*Fis*] = -1 / (2*N* -1); Robertson, 1965). A two-tailed t-test was run to compare observed locus *Fis* values against E[*Fis*] to classify populations into three categories: 1) small-*N* sampling bias (p > 0.05), where observed *Fis* does not deviate from theoretical expectations; 2) balancing selection (p < 0.05, *Fis* < E[*Fis*]), where negative *Fis* exceeds small-*N* expectations; and 3) inbreeding/selfing (p < 0.05, *Fis* > E[*Fis*]), where loci are more homozygous than expected under small-*N* expectations. Finally, individual multilocus heterozygosity was calculated using inbreedR v. 0.3.3 (Stoffel et al., 2016) to assess genome-wide heterozygosity across individuals and populations for the gid_bent_cleaned dataset.

Genome-wide identity disequilibrium was quantified for *C. bentleyi* using inbreedR to evaluate whether observed patterns of excess heterozygosity reflect a genome-wide phenomenon versus isolated locus-specific effects. The *g_2_* statistic was estimated from a binary heterozygosity matrix across SNP loci for the gid_bentleyi dataset using 1,000 bootstrap pseudoreplicates for confidence interval estimation and 1,000 permutations to test for significance against the null hypothesis of no identity disequilibrium (David et al., 2007; Stoffel et al., 2016). To assess the accuracy with which individual multilocus heterozygosity reflects genome-wide prevalence, we calculated the heterozygosity-heterozygosity correlation (*rhh*; Balloux et al., 2004) across randomly split halves of the data using 100 iterations in inbreedR.

### 2.8. Tests for bottlenecks and demographic history modeling

To test for recent bottleneck signatures, we evaluated population-level heterozygosity excess across 2,347 linkage-disequilibrium-thinned loci (gid_bentleyi) using the Cornuet and Luikart (1996) framework. This test quantifies whether *H_O_* is greater than the *H_E_* expected under mutation-drift equilibrium. We performed a paired, two-tailed Wilcoxon signed-rank test comparing *H_O_* against the *H_E_* mutation-drift equilibrium expectations derived from observed allele counts per locus. We then evaluated whether individual multilocus heterozygosity (*MLH*) across 52,718 unpruned loci (gid_bent_unpruned) significantly exceeded Hardy-Weinberg Equilibrium (HWE) expectations. We extracted locus-specific allele frequencies to model null HWE genotype distributions and generated 1,000 simulated null populations by drawing genotypes from a binomial distribution. The *p*-values were calculated as the proportion of simulated HWE iterations where the simulated population mean *MLH* equaled or exceeded the observed population mean *MLH*.

To evaluate demographic history and test for population bottlenecks, we performed Approximate Bayesian Computation (ABC) across six demographic scenarios including divergence timing [Pre-Last Glacial Maximum (Pre-LGM; 20-12 kya) vs. Post-LGM; 100-12 kya], post-glacial dynamics (expansion vs. bottleneck), and 200-year anthropogenic disturbance dynamics (recent bottleneck vs. no bottleneck). Analyses were conducted with the R package abc v.2.2.2 (Csilléry et al., 2012). Model selection was conducted by comparing an empirical vector to 10^6^ coalescent simulations per model evaluated across a six-metric summary statistic vector consisting of mean *H_O_*, mean *H_E_*, *ΔH* (*H_O_* - *H_E_*), mean *F_ST_*, variance of *F_ST_*, and multilocus linkage disequilibrium (*r̄_D_*). Posterior probabilities of the models were estimated with multinomial logistic regression (tolerance = 0.02). To evaluate the classification success and error rates of our models, we performed cross-validation across 100 pseudo-observed datasets per model.

### 2.9. Simulations under alternative selection scenarios

To investigate patterns expected under alternative selection mechanisms maintaining excess heterozygosity and negative inbreeding coefficients, we performed simulations testing scenarios of neutrality, overdominance, and negative frequency dependent selection (NFDS) in R. We compared our observed *F_IS_* distribution with 28 parameterized models, varying selfing rates, population sizes, overdominance selection intensity, and negative frequency-dependent selection intensity. Selfing rates were specified as either 100% (obligate selfing) or 90% (predominant selfing). Population sizes were specified as *N* = 10 or *N* = 100, in accordance with observations in the field. Overdominance parameters were specified as relative fitnesses of each genotype at each locus (*w*) as follows: neutral (*w_homozygote_* = *w_heterozygote_* = 1), weak (*w_homozygote_* = 0.85), moderate (*w_homozygote_* = 0.5), strong (*w_homozygote_* = 0.15), and lethal (*w_homozygote_* = 0.0). NFDS was simulated by scaling relative genotype fitness inversely to population frequency, where common genotypes experienced reduced fitness according to a selection coefficient of 0.3 for weak selection and 0.8 for strong selection. Genotype fitness values were bound at a minimum of 0.01 to prevent negative probabilities during parental sampling. Parallel simulations were computed in R using the furrr package v.0.3.1 (Vaughan and Dancho, 2022). Each parameter combination was run across 500 loci for 50 generations, tracking genotype frequencies to calculate final locus-specific inbreeding coefficients. Goodness-of-fit between simulated and empirical inbreeding coefficient distributions was summarized by Wasserstein Distances (*D_W_*), i.e., the minimum cost of transforming the observed *F_IS_* distribution into each simulated distribution (Vaseršteĭn, 1969; Dobrushin, 1970; Panaretos and Zemel, 2018) using the transport package v.0.15 (Schumacher et al., 2024). The likelihood of each simulated model is given by *L_model_* = exp((-1/2)(*D_W-model_*/*ε*)^2^), where *ε* = the minimum *D_W_* among each of the simulated model distributions and the empirical *F_IS_* distribution, and the weight of each model is given by *W_model_* = *L_model_* / Σ(*L_all models_*).

## 3. Results

### 3.1. *Corallorhiza bentleyi* individuals are predominantly cleistogamous or mixed chasmogamous-cleistogamous

Surveys at six sites over four years detected no racemes at the OR Giles VA and WR Giles VA sites (visited in 2022 and 2024). The two Bath County, Virginia sites were visited only in 2022, yielding three racemes at CG Bath VA (27 chamogamous/13 cleistogamous flowers, hereafter 27/13), while site DK Bath VA yielded only two racemes (2/14). The PM Monroe County, West Virginia site (type locality) was visited in 2022 and 2024, yielding a total of nine individuals in 2022 (11/85) and 13 individuals in 2024 (15/152). Visits to the PC Pocahontas WV site in 2022, 2024, and 2025 yielded 12 (29/77), 10 (1/129), and 13 (29/184) individuals, respectively. Plants at the Bath County, Virginia sites were predominantly chasmogamous (CG) or mixed (DK), though only five individuals were observed among these two localities (Fig. 2). Based on data from the two West Virginia sites, in which most of the observations were made, individual flowers were predominantly cleistogamous, and individual plants were predominantly fully cleistogamous or mixed (Fig. 2).

**Figure 2.**
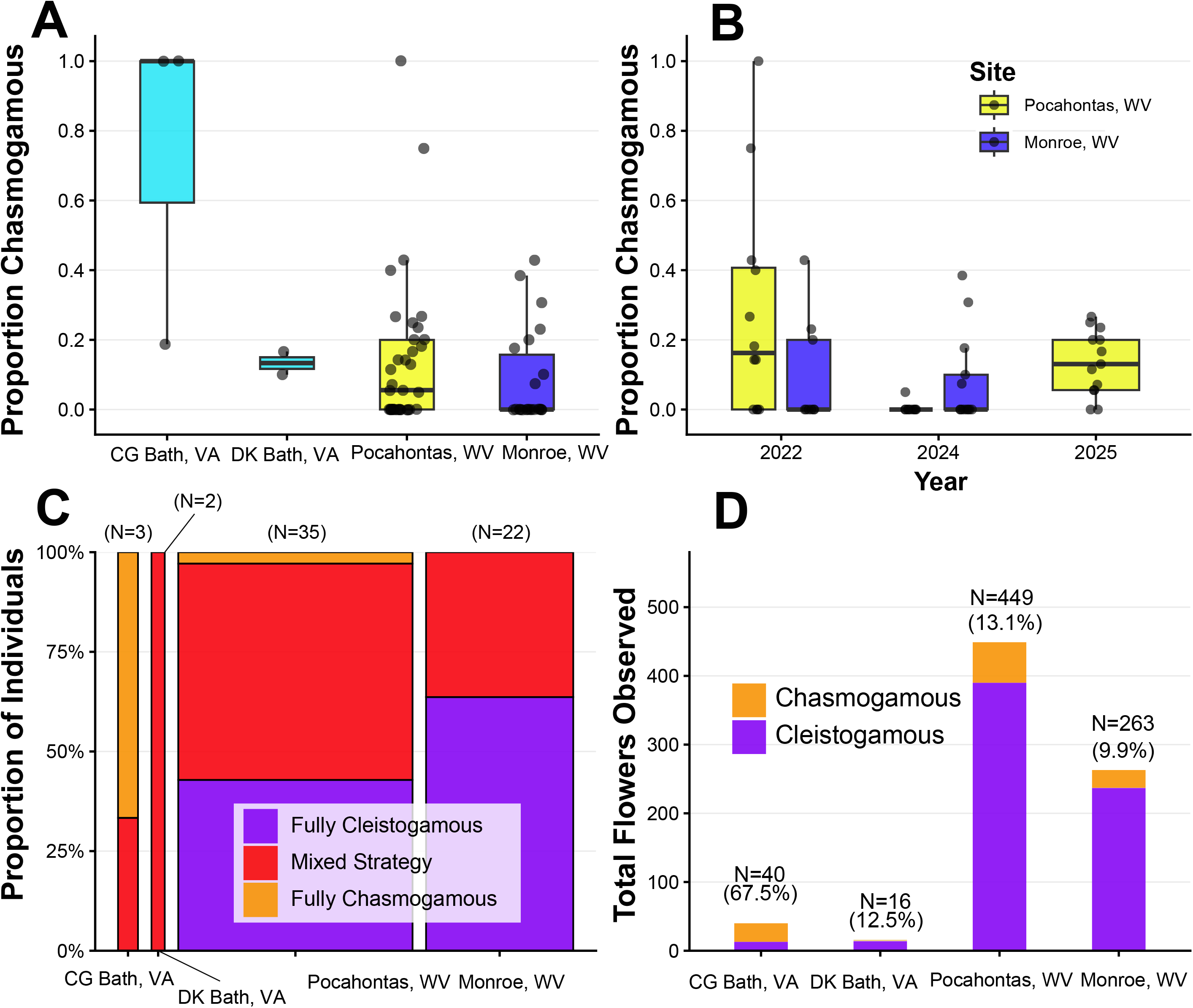
**A**. total proportion of chasmogamous (open) flowers observed among four localities in *Corallorhiza bentleyi* (hereafter, from 2022-2025; no observation efforts were made in 2023). **B**. Proportions of chasmogamous flowers observed across three observation years for two sites (Monroe County, West Virginia, USA and Pocahontas County, West Virginia, USA). **C**. Total proportions of individual plants that were observed to be fully cleistogamous (closed-flowered), fully chasmogamous, or mixed. **D**. Total numbers of flowers observed as either chasmogamous or cleistogamous.

Examination of physical and digitized specimens of *Corallorhiza involuta* yielded only open-flowered individuals. The range of floral “openness” varied widely, however, from individuals bearing flowers with all perianth parts clearly separated to those with only the lip protruding. Of 14 iNaturalist records that we could positively identify, two were determined to be nearly cleistogamous. Both were at the periphery of the known range of *C. involuta*, in the states of Chiapas (https://www.inaturalist.org/observations/357023250) and Michoacán (https://www.inaturalist.org/observations/4293604). Examination of representative liquid-preserved flowers of both *C. bentleyi* and *C. involuta* revealed that the pollinia were swollen in-place (i.e., the pollen grains had germinated) and had contacted the stigma, suggesting a mechanism for autopollination (Fig. 1).

### 3.2. *Corallorhiza involuta* and *C. bentleyi* are distinct, with weak population structure in *C. bentleyi*

Nucleotide diversity (π) within *C. involuta* was 0.2219 (n = 5), within *C. bentleyi* was 0.1928 ± 0.0156 (5 randomly sampled individuals, repeated 10×), and between the two species was 0.2495 ± 0.0071. Maximum likelihood phylogenetic analysis of 2,347 filtered SNPs revealed two principal clades, corresponding to *C. involuta* and *C. bentleyi* (Fig. 3A). Each clade received 100% bootstrap support (BS, hereafter) and was separated by relatively long branches, indicating a pattern of reciprocal exclusivity among accession of both taxa. Accessions of *C. involuta* grouped by sampling locality (BS = 100% for Mex-MX from the state of Mexico and 98% for Mor-MX from the state of Morelos). Within *C. bentleyi*, all accessions from Bath County, Virginia grouped together (BS = 99%), as did accessions from Monroe County, West Virginia (BS = 98%). Accessions from all other localities, however, clustered within mixed-locality groupings. UPGMA analysis among localities similarly showed a strongly supported sister relationship among *C. bentleyi* and *C. involuta* (BS = 100%; Fig. 3B). Within *C. bentleyi*, locality PM Monroe WV clustered as sister of all other localities, followed by WR Giles VA, OR Giles VA, RG Alleghany VA, and CG Bath VA. Internal branch distances indicated low divergence among localities.

**Figure 3.**
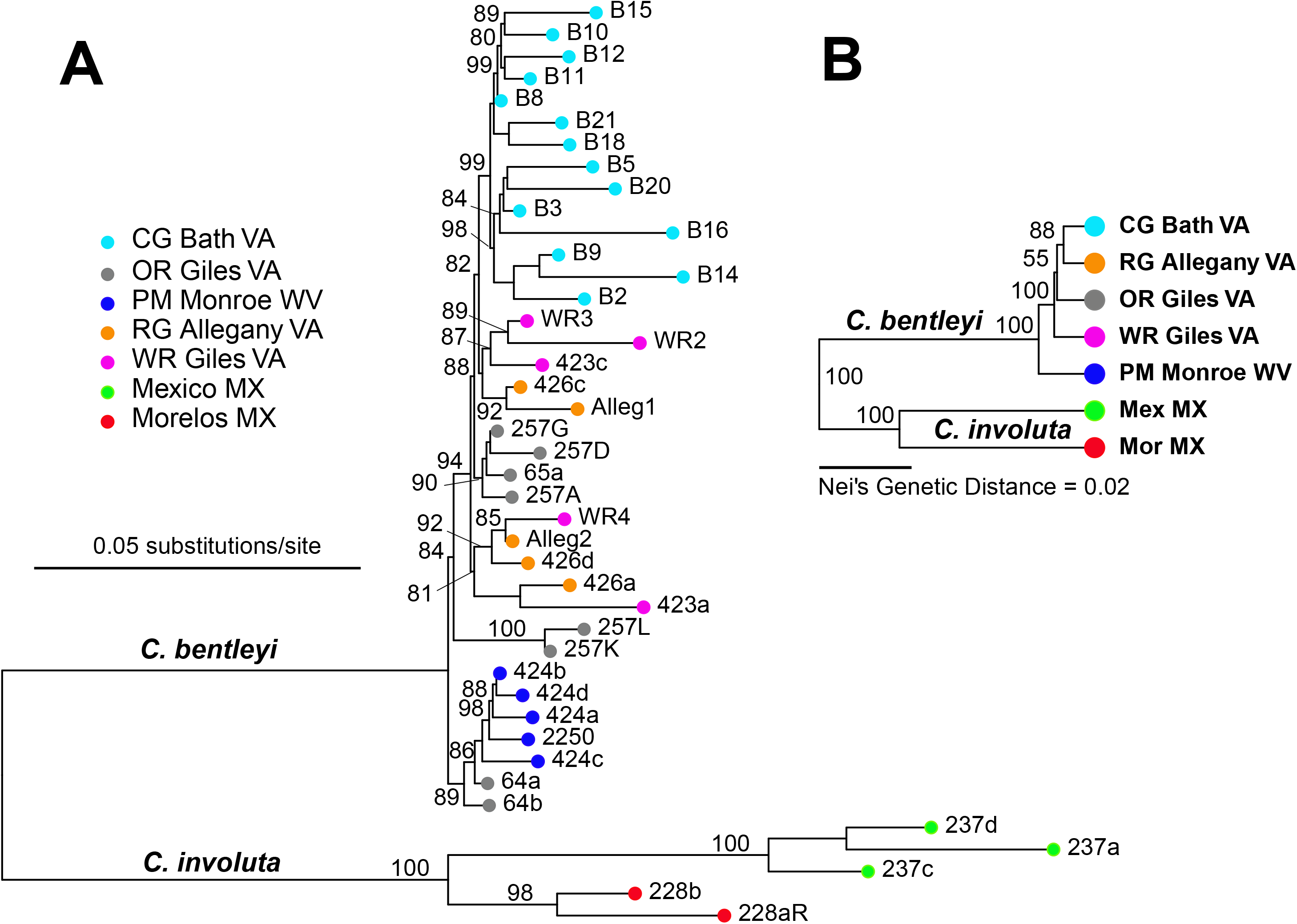
**A**. Phylogenetic tree based on maximum likelihood analysis of 2,347 biallelic SNPs among *Corallorhiza involuta* and *C. bentleyi* under the TVM+F+ASC+R3+ASC model in IQTREE3 (BIC = 84858.2, BIC weight = 0.95). Numbers adjacent to branches are based on 1,000 ultra-fast bootstrap pseudoreplicates. Scale bar = 0.05 substitutions/site. **B**. UPGMA clustering based on Nei’s Genetic distance among sampling localities of *C. involuta* and *C. bentleyi*. Scale bar: Nei’s Genetic Distance = 0.02. Numbers adjacent to branches are based on 1,000 bootstrap pseudoreplicates conducted with the R package poppr.

Principal components analysis demonstrated clear separation of *C. involuta* from *C. bentleyi* along PC axis 1, and further between individuals of *C. involuta* from the two Mexican localities along PC axis 2 (Fig. 4A). PC axis 3 also separated the two localities of *C. involuta* from each other and *C. bentleyi*. Analysis of the *C. bentleyi*-only dataset (gid_bentleyi) revealed a weak pattern of separation among the sampled localities (Fig. 4C). PC axis 1 partially separated individuals from OR Giles VA from CG Bath VA, and RG Alleghany VA, and WR Giles VA. No clear patterns of separation among localities were evident along PC axes 3 or 4 (Fig. 4D).

**Figure 4.**
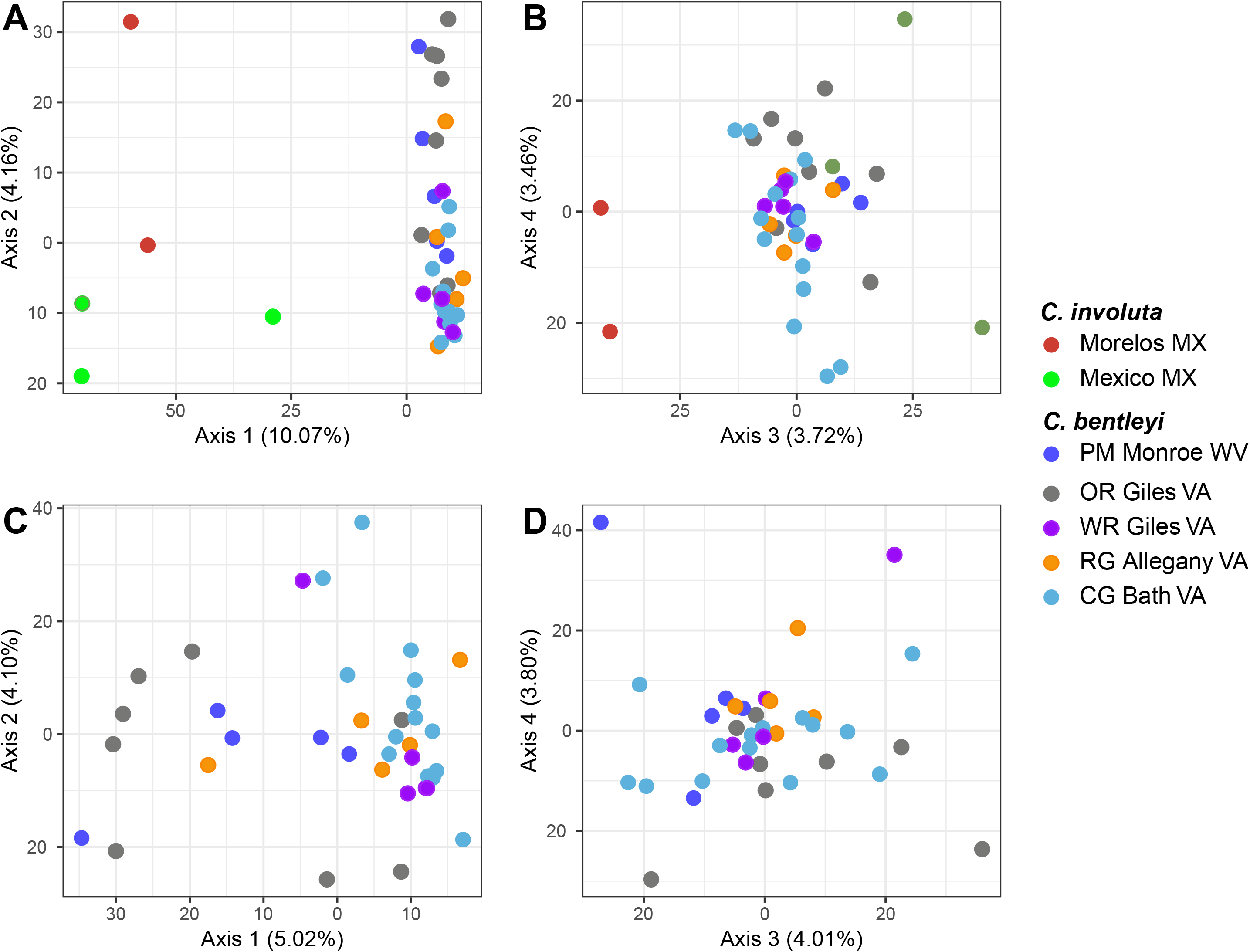
**A**. Principal components analysis (PCA) of 2,347 unlinked SNPs for the full dataset (*C. involuta* and *C. bentleyi*) showing PC axes 1 and 2, and **B**. axes 3 and 4. Percentages are the total amount of variation explained by each axis. **C**. Principal components analysis of unlinked SNPs for the *C. bentleyi*-only dataset showing PC axes 1 and 2, and **D**. axes 3 and 4.

Results from unsupervised population structure analyses revealed that the optimal number of ancestral genomic clusters for the *C. bentleyi*-only dataset was *K* = 1 (based on the lowest cross-entropy value in the sNMF analysis and the lowest BIC score in the DAPC analysis; Fig. S1).

Results from the supervised DAPC analysis, with sampling localities set as population priors (optimal number of PCs = 6, max A-score = 0.42), revealed finer-scale population-specific signatures of differentiation (Fig. 5A). All individuals from PM Monroe WV (blue) and OR Giles VA (grey) were assigned to their home populations with high posterior membership probabilities (> 0.90%; Fig. 5A). Individuals from CG Bath VA (cyan) were predominantly assigned to their native cluster, though several individuals exhibited modest patterns of mixed ancestry. By contrast, WR Giles VA (purple) and RG Allegany VA (orange) demonstrated substantial mixed ancestry, sharing proportions of membership across multiple clusters (Fig. 5A). Ordination across the first two Linear Discriminant axes (Fig. 5B) clearly separated PM Monroe WV along Linear Discriminant 1 (LD1), forming a distinct, non-overlapping cluster relative to all Virginia localities. Axis LD2 further separated CG Bath VA along the upper vertical axis (LD2), while OR Giles VA separated toward the lower-right quadrant. Extensive overlap was observed among WR Giles VA, RG Allegany VA, and CG Bath VA.

**Figure 5.**
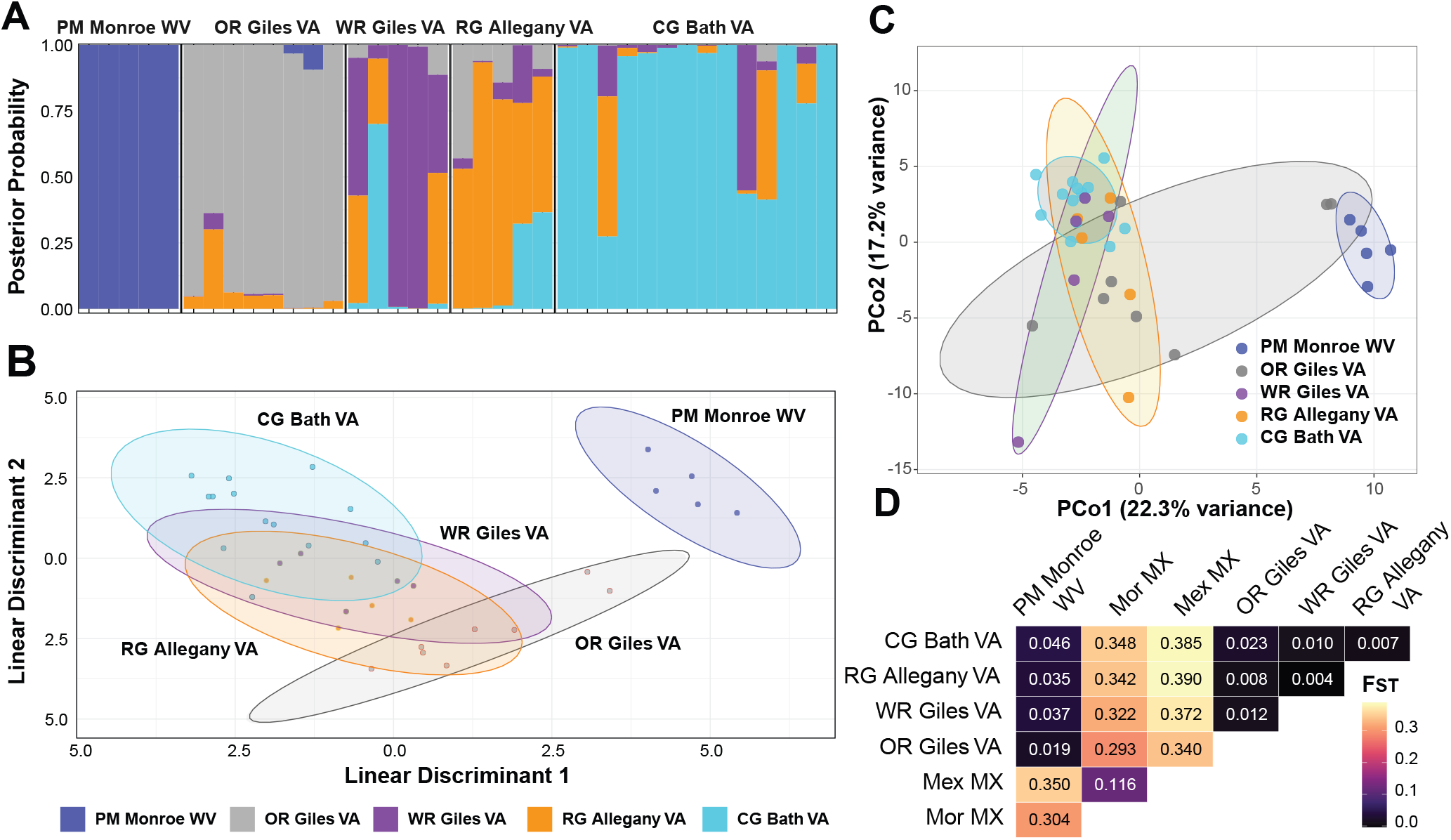
**A.** Supervised discriminant analysis of principal components (DAPC) for the *C. bentleyi*-only dataset, with sampling locality treated as the population prior, showing barplot of inferred DAPC ancestry coefficients. **B**. Scatterplot of the first two linear discriminant axes for the supervised DAPC. Legend colors correspond to both A and B. **C**. Principal coordinates analysis (PCoA) of the coancestry matrix for *C. bentleyi* [*D = max(Coancestry) – Coancestry*] from analysis with fineRADstructure. **D**. Pairwise differentiation (*F_ST_*) among sampling localities for *C. involuta* and *C. bentleyi*.

Principal coordinates analysis on the fineRADstructure coancestry matrix (Fig. 5C) mirrored the patterns observed in the supervised DAPC analysis (Fig. 5B). PM Monroe WV (blue) formed a cluster separated from all other localities along PCo axis 1. Similarly, CG Bath VA (cyan) formed a cluster concentrated in the upper-left quadrant. However, two OR Giles VA individuals (accessions 64a and 64b) clustered closely with PM Monroe WV, while the remaining OR individuals occupied a cluster of Virginia localities (Fig. 5C). Pairwise *F_ST_* analysis demonstrated differentiation among *C. bentleyi* and *C. involuta* (*F_ST_* > 0.29 in all comparisons), and to a lesser extent, between the two *C. involuta* localities (*F_ST_* = 0.116; Fig. 5D). Within *C. bentleyi*, *F_ST_* values were low, from 0.004 (RG Alleghany VA vs. Giles, VA) to 0.046 (PM Monroe WV vs. CG Bath VA). A Mantel test revealed weak but non-significant isolation by distance (Mantel *r*^2^ = 0.517, p = 0.108). Analysis of Molecular Variance (Table 2) revealed most variation was explained between species (47.47%, *p* = 0.042) and within localities (49.87%, *p* < 0.001), while little variation was explained between localities within species (2.66%, *p* < 0.001).

**Table 2.**
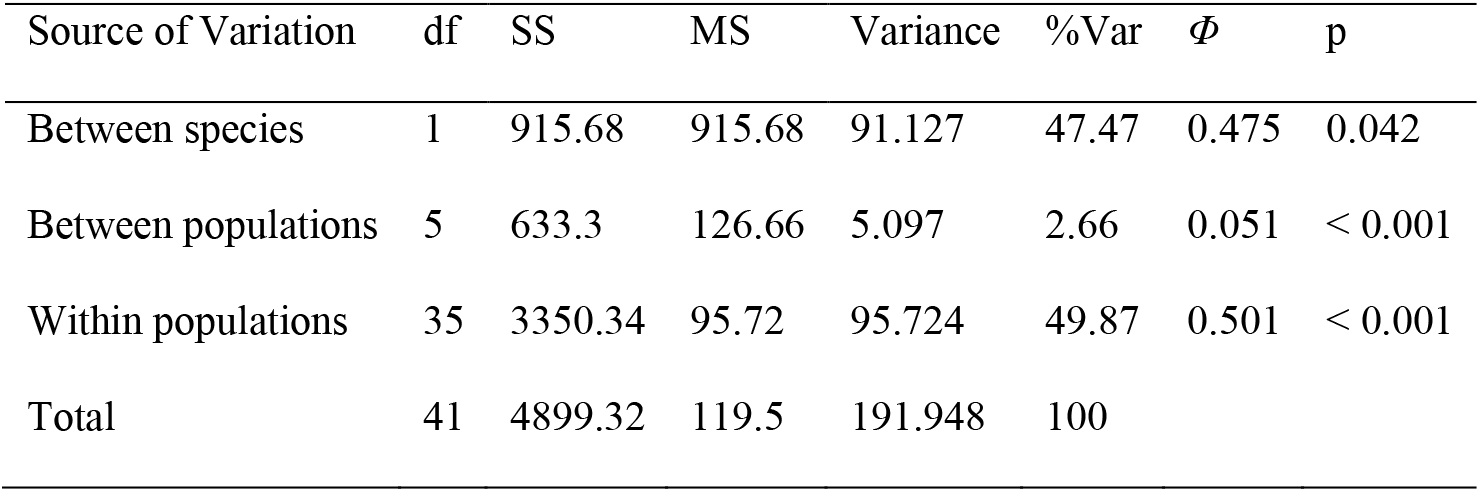
Hierarchical analysis of molecular variance (AMOVA). ‘df’ = degrees of freedom, ‘SS’ = sum of square, ‘MS’ = mean squares, ‘Variance’ = the variance component of each level, ‘%Var’ = the percent of total variation explained by each level, ‘*Φ*’= the phi statistic for each level, and ‘p’ = the p-value based on 999 randomizations for the significance of each variance component.

| Source of Variation | df | SS | MS | Variance | %Var | $\Phi$ | p |
| --- | --- | --- | --- | --- | --- | --- | --- |
| Between species | 1 | 915.68 | 915.68 | 91.127 | 47.47 | 0.475 | 0.042 |
| Between populations | 5 | 633.3 | 126.66 | 5.097 | 2.66 | 0.051 | < 0.001 |
| Within populations | 35 | 3350.34 | 95.72 | 95.724 | 49.87 | 0.501 | < 0.001 |
| Total | 41 | 4899.32 | 119.5 | 191.948 | 100 |  |  |

### 3.3. *Corallorhiza bentleyi* displays unexpected patterns of excess heterozygosity and negative inbreeding coefficients

Overall, values of observed heterozygosity (*H_O_*), expected heterozygosity *H_E_*, *H_O_*/*H_E_* ratio, and inbreeding coefficient (*F_IS_*) were similar across localities of both *C. involuta* and *C. bentleyi* (Table 3). Mean *H_O_* for each locality of *C. bentleyi* and *C. involuta* ranged from 0.141 (*C. involuta* Mex-MX) to 0.201 (PM Monroe WV) and *H_E_* ranged from 0.120 (*C. involuta* Mex-MX) to 0.170 (OR Giles VA). Surprisingly, an excess of heterozygosity (Mean *H_O_*/*H_E_*) was observed in all sampling localities for both species, ranging from 1.134 (OR Giles VA) to 1.418 (*C. involuta* Mor-MX). Inbreeding coefficients were negative across all localities in both species, with the lowest being -0.418 for *C. involuta* Mor-MX and the highest being -0.120 for *C. bentleyi* CG Bath VA. Numbers of private alleles reflected sample size variation, though the relationship was not significant (Spearman’s rank correlation *S* = 66.377, *p* = 0.691). Within *C. bentleyi*, the more extensive sampling at CG-Bath-VA yielded 193 private alleles, whereas localities with smaller sample sizes harbored fewer unique variants. Within *C. involuta*, Mor-MX yielded 68 private alleles while Mex-MX yielded 305. Analysis of linkage disequilibrium revealed significant but low overall multilocus association across populations (*I_A_* = 48.905, *r̄*_D_ = 0.00922, *p* = 0.005). Among individual sampling localities, all *C. bentleyi* populations but PM Monroe WV exhibited low but statistically significant linkage disequilibrium (*p* ≤ 0.01), with the highest standardized index observed in Mex-MX (*r̄*_D_ = 0.08877).

**Table 3.**
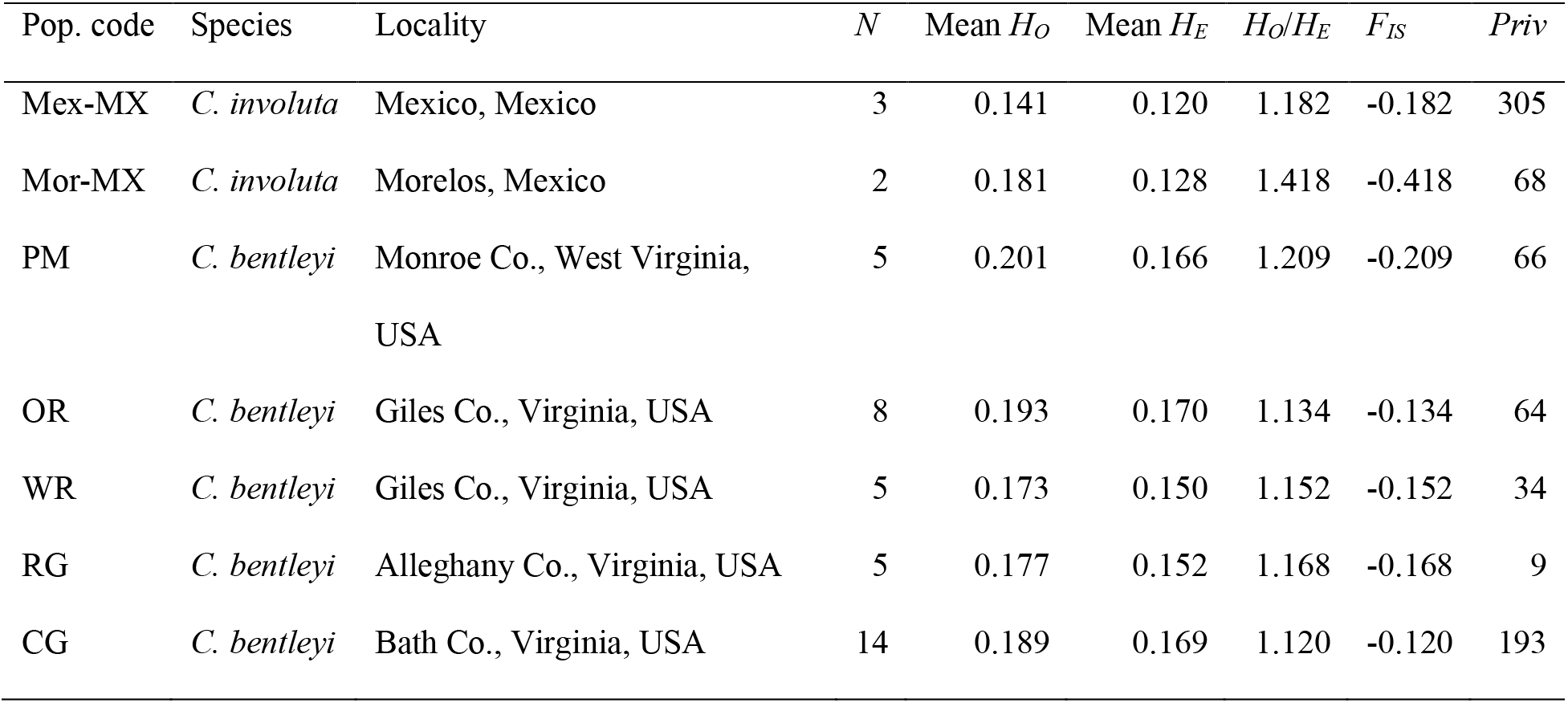
Summary of population genomic diversity for each sampling locality of *Corallorhiza involuta* and *C*. *bentleyi*. ‘Pop. code’ = population code for each sampling locality; N = number of individuals sampled; Mean *H_O_* = mean observed heterozygosity across all loci; Mean *H_E_* = mean expected heterozygosity across all loci; *H_O_*/*H_E_* mean ratio of observed to expected heterozygosity; *F_IS_* = inbreeding coefficient; *Priv* = number of private alleles.

| Pop. code | Species | Locality | N | Mean $H_O$ | Mean $H_E$ | $H_O/H_E$ | $F_{IS}$ | $Priv$ |
| --- | --- | --- | --- | --- | --- | --- | --- | --- |
| Mex-MX | *C. involuta* | Mexico, Mexico | 3 | 0.141 | 0.120 | 1.182 | -0.182 | 305 |
| Mor-MX | *C. involuta* | Morelos, Mexico | 2 | 0.181 | 0.128 | 1.418 | -0.418 | 68 |
| PM | *C. bentleyi* | Monroe Co., West Virginia,<br>USA | 5 | 0.201 | 0.166 | 1.209 | -0.209 | 66 |
| OR | *C. bentleyi* | Giles Co., Virginia, USA | 8 | 0.193 | 0.170 | 1.134 | -0.134 | 64 |
| WR | *C. bentleyi* | Giles Co., Virginia, USA | 5 | 0.173 | 0.150 | 1.152 | -0.152 | 34 |
| RG | *C. bentleyi* | Alleghany Co., Virginia, USA | 5 | 0.177 | 0.152 | 1.168 | -0.168 | 9 |
| CG | *C. bentleyi* | Bath Co., Virginia, USA | 14 | 0.189 | 0.169 | 1.120 | -0.120 | 193 |

### 3.4. Excess heterozygosity and negative inbreeding are not explained by bioinformatic artifacts or small-*N* sampling

Comparison of our original heterozygosity and inbreeding coefficients with those from the discoSNP reference-free assembly yielded similar results overall, with *H_O_*/*H_E_* > 1 and *F_IS_* < 0 for all localities (Fig. S2). Genome-wide assessments of locus-level diversity revealed a strongly right-skewed distribution of *H_O_* (Fig. 6A) and a significant overall excess of heterozygosity across loci (*ΔH* = +0.038, *p* < 0.0001; Fig. 6B). Furthermore, minor allele frequency (*MAF*) displayed a negative relationship with inbreeding coefficients across loci (*F_IS_*), where higher-*MAF* loci exhibited reduced *F_IS_* values under both the non-pruned data (*β* = -1.61, *p* < 0.0001; gid_bent_unpruned) and high-*H_O_*-pruned data (*β* = -1.27, *p* < 0.0001; gid_bent_cleaned; Fig. 6C). At the population level, observed *F_IS_* estimates across all five sampling localities were significantly negative and deviated below expectations under finite sample-size bias alone (*p* < 0.001 in all cases; Fig. 6D). Across individuals, multilocus heterozygosity (*MLH*) was consistently high, with median values per site between 0.43 and 0.50, alongside occasional low-*MLH* outliers reaching as low as 0.31 (Fig. 6E). Finally, significant genome-wide identity disequilibrium was detected across the 37 *C. bentleyi* individuals (*g_2_* = 0.0623, *p* = 0.001). Mean heterozygosity-heterozygosity correlation was high (*r_hh_* = 0.9495), indicating that half of all randomly selected SNPs captured 95 percent of the variance in individual genome-wide heterozygosity in the other half.

**Figure 6.**
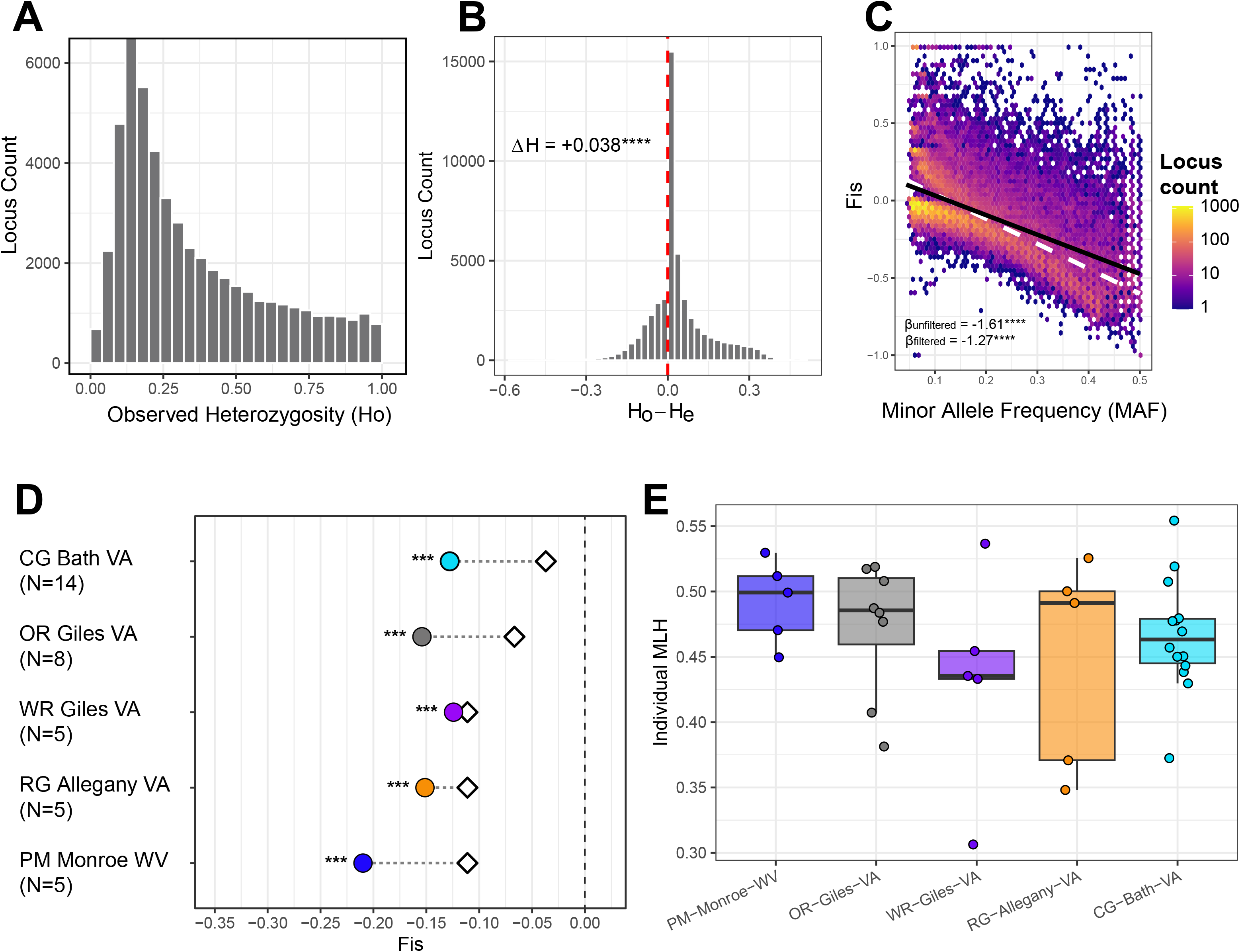
Patterns in heterozygosity and inbreeding coefficients in *C. bentleyi*. **A**. Frequency of loci as a function of observed heterozygosity (*H_O_*) for the full, non-pruned *Corallorhiza bentleyi* data (gid_bent_unpruned; see Table 1). **B**. Analysis of excess heterozygosity after removing sites with *H_O_* > 0.85 (gid_bent_cleaned dataset). *ΔH* = *H_O_* – *H_E_* for each individual locus, where *ΔH* ∼ 0 is the null expectation of no excess heterozygosity and *ΔH* > 0 indicates excess heterozygosity. *p*-value is based on a one-tailed t-test of observed *ΔH* vs. the null expectation of *ΔH* = 0 (**** *p* < 0.0001). **C**. Plot of minor allele frequencies vs. inbreeding coefficients before (gid_bent_unpruned, black solid line) and after removal of sites with *H_O_* > 0.85 (gid_bent_cleaned, white dashed line). *β* values = the slope of the lines, and **** *p* < 0.0001. **D**. Observed *F_IS_* values (colored circles) vs *F_IS_* under the small-*N* expectation (empty diamonds), defined as E[*F_IS_*] = -1 / (2*N* -1) (Robertson, 1965; Hill and Robertson 1966). *** *p* < 0.001. **E**. Individual multilocus heterozygosity (*MLH*).

### 3.5. Demographic modeling rejects bottlenecks despite widespread heterozygosity excess

Population-level bottleneck tests (Cornuet and Luikart, 1996) across 2,347 LD-thinned loci revealed significant excess *H_O_* relative to expectations across all localities (p < 0.0001; Table 5). Across 52,718 unpruned loci, observed multilocus heterozygosity was significantly elevated above random-mating HWE expectations across all *Corallorhiza bentleyi* localities (p < 0.0001; Table 5). Observed *MLH* ranged from 0.2842 in WR-Giles-VA to 0.3427 in PM-Monroe-WV, compared to HWE expectations ranging from 0.2366 to 0.2582. However, demographic model selection via Approximate Bayesian Computation (ABC) demonstrated that this excess heterozygosity is not associated with population bottlenecks: model comparison yielded the highest support for a post-glacial range expansion model without recent bottlenecks (Model 1, *PP* = 1.0, Table 6). All alternative models, including those incorporating recent anthropogenic bottlenecks or pre-LGM divergence, were rejected (*PP* = 0, *BF* = ∞). Cross-validation demonstrated high classification power, with several models including the one with the lowest-BIC (Model 1, post-LGM divergence and no bottlenecks) achieving classification accuracies of 100%.

**Table 4.** Analysis of linkage disequilibrium for each sampling locality of *Corallorhiza involuta* and *C. bentleyi*. *I_A_* = index of association; p(*I_A_*) = significance of the index of association based on 199 randomizations; *r̄*_D_ = standardized index of association; p(*r̄*_D_) = significance of the standardized index of association based on 199 randomizations. ‘*NA’* values for Mor-Mex: linkage metrics were not calculated due to low sample size (*N* = 2).

| Pop. code | Species | $I_A$ | $p(I_A)$ | $\bar{r}_D$ | $p(\bar{r}_D)$ |
| --- | --- | --- | --- | --- | --- |
| Overall |  | 48.905 | 0.005 | 0.00922 | 0.005 |
| Mex-MX | *C. bentleyi* | 142.359 | 0.01 | 0.08877 | 0.01 |
| Mor-MX | *C. involuta* | NA | NA | NA | NA |
| PM | *C. involuta* | 0.834 | 0.1 | 0.00025 | 0.105 |
| OR | *C. bentleyi* | 7.304 | 0.005 | 0.00182 | 0.005 |
| WR | *C. bentleyi* | 22.346 | 0.005 | 0.00731 | 0.005 |
| RG | *C. bentleyi* | 27.662 | 0.005 | 0.00893 | 0.005 |
| CG | *C. bentleyi* | 5.174 | 0.005 | 0.00113 | 0.005 |

**Table 5.** Tests of population bottlenecks and putative overdominance. Bottleneck analysis (gid_bentleyi data, 2,347 SNP loci): *N* = sample size per locality, *H_O_* (mean) = mean observed heterozygosity, *H_E_* (mean) = mean expected heterozygosity under mutation-drift equilibrium (Cornuet et al., 1996), *A* = mean number of alleles per locus, Wilcoxon *V* = sum-of-positive-signed-ranks statistic, *p* = significance of two-tailed Wilcoxon signed rank test. Overdominance (gid_bentleyi_unpruned; 52,718 SNP loci): *MLH_O_* = mean multilocus observed heterozygosity; *MLH_E_* = mean multilocus expected heterozygosity, p = proportion of 1,000 simulated HWE iterations where the simulated population mean *MLH* equaled or exceeded the observed population mean *MLH*.

| Population | $N$ | $H_O$ (mean) | $H_E$ (mean) | $A$ | Wilcoxon $V$ | $p$ | $MLH_O$ | $MLH_E$ | $p$ |
| --- | --- | --- | --- | --- | --- | --- | --- | --- | --- |
| PM-Monroe-WV | 5 | 0.2414 | 0.1845 | 1.64 | 1070005 | < 0.0001 | 0.4919 | 0.2547 | < 0.0001 |
| OR-Giles-VA | 8 | 0.2446 | 0.1975 | 1.78 | 1538777 | < 0.0001 | 0.4722 | 0.2574 | < 0.0001 |
| WR-Giles-VA | 5 | 0.1848 | 0.1601 | 1.55 | 708238.5 | < 0.0001 | 0.4322 | 0.2366 | < 0.0001 |
| RG-Allegany-VA | 5 | 0.1982 | 0.1583 | 1.56 | 805829.5 | < 0.0001 | 0.4463 | 0.241 | < 0.0001 |
| CG-Bath-VA | 14 | 0.1927 | 0.1694 | 1.83 | 1592848 | < 0.0001 | 0.4658 | 0.2582 | < 0.0001 |

**Table 6.** Results from demographic model testing in the R package abc among six demographic models. Divergence: ‘Post-LGM’ = after the Last Glacial Maximum (< 12 kya), ‘Pre-LGM’ = before the Last Glacial Maximum (∼100 kya). LGM and Recent Dynamics indicate inclusion of bottleneck/no bottleneck 10-12 kya or 100-200 ya. *P* = posterior probability of the model, *BF* = Bayes Factor of each model compared to the highest posterior probability model, and *CV* = the proportion of successful cross-validation tests.

| Model | Divergence | LGM Dynamics | Recent Dynamics | $P$ | $BF$ | $CV$ |
| --- | --- | --- | --- | --- | --- | --- |
| 1 | Post-LGM | Expansion (Both) | No bottleneck | 1 | 1 | 1 |
| 2 | Post-LGM | Bottleneck (*C. bentleyi*) | Bottleneck (*C. bentleyi*) | 0 | Inf | 0.94 |
| 3 | Post-LGM | Bottleneck (*C. bentleyi*) | No bottleneck | 0 | Inf | 1 |
| 4 | Post-LGM | Expansion (Both) | Bottleneck (*C. bentleyi*) | 0 | Inf | 0.83 |
| 5 | Post-LGM | Expansion (Both) | Bottleneck (Both) | 0 | Inf | 0.58 |
| 6 | Pre-LGM | Expansion (Both) | Bottleneck (*C. bentleyi*) | 0 | Inf | 1 |

### 3.6. Simulations support overdominance over NFDS

Out of the 28 simulation runs, the 11 top-scoring models all included overdominance rather than negative frequency dependent selection (NFDS; Fig. 7; Table S1). The overall best-fit model specified overdominance, *N* = 100, selfing rate = 100%, and *w_homozygote_* = 0 (i.e., homozygote fitness = “lethal”), having the lowest Wasserstein Distance to the empirical *F_IS_* distribution as well as the highest model weight (*W-distance* = 0.1119, weight = 0.1937; Fig. 7; Table S1). The next three models all included overdominance and “lethal” *w_homozygote_*, differing only in *N* and selfing rates (*W-distances* = 0.1651-0.1189, weights = 0.1491-0.1882; Table S1). By contrast, the top-scoring NFDS model was 12th overall when ranked among all 28 models (*N* = 10, selfing rate = 90%, selection intensity = 0.8, *W-distance* = 0.7991, weight = 2.87×10^−6^; Fig. 7; Table S1).

**Figure 7.**
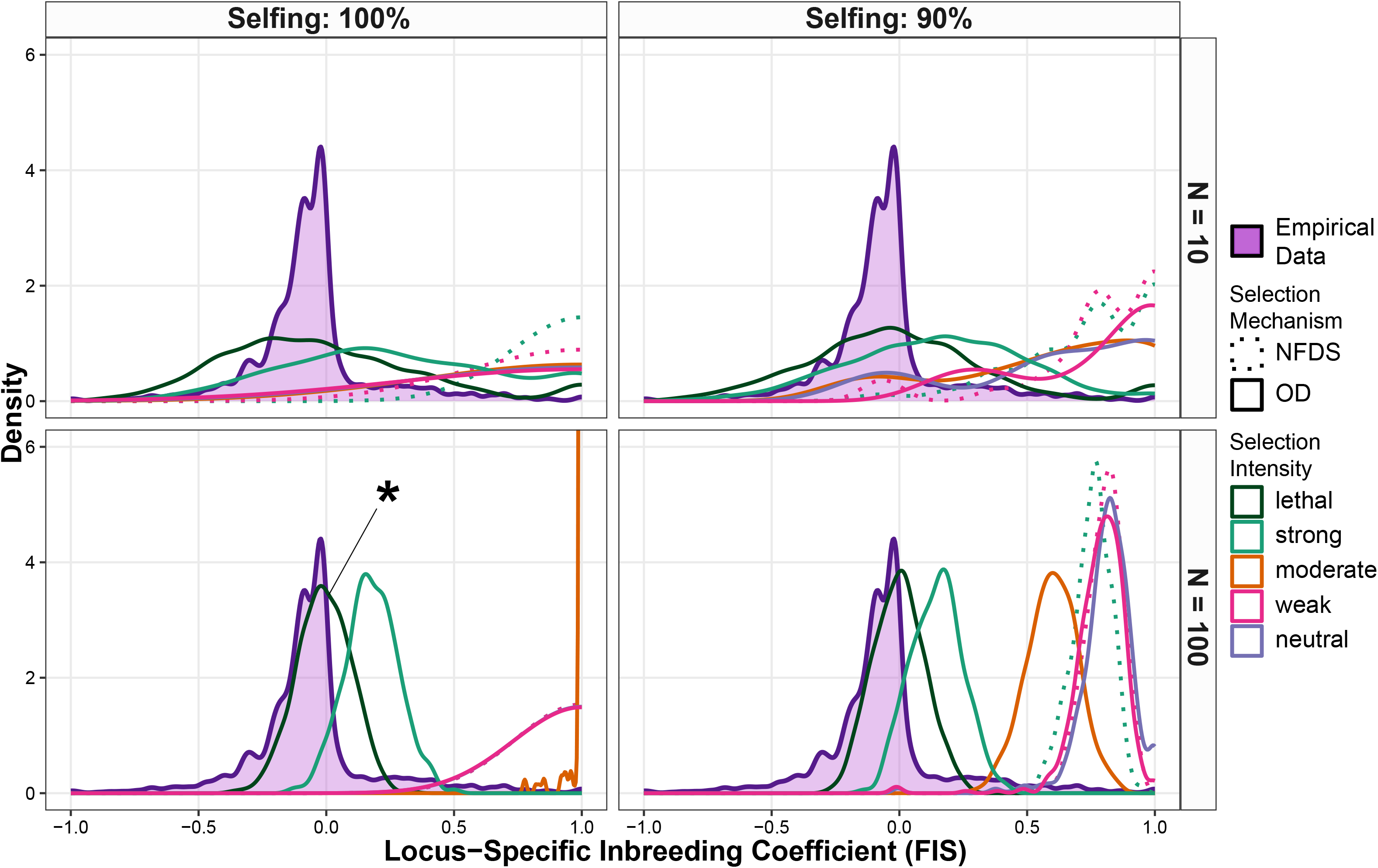
Density plots from simulations of *F_IS_* distributions (inbreeding coefficient) under 28 model parameter sets compared to the empirical *F_IS_* distribution for overdominance (OD) and negative frequency dependent selection (NFDS). Overdominance parameters (solid lines) for relative fitnesses (*w*): neutral (*w_homozygote_* = *w_heterozygote_* = 1, purple), weak (*w_homozygote_* = 0.85, magenta), moderate (*w_homozygote_* = 0.5, orange), strong (*w_homozygote_* = 0.15, aqua), and lethal (*w_homozygote_* = 0.0, green). NFDS parameters (dashed lines): weak (*w* = 0.3, magenta); strong (*w* = 0.8, aqua). The empirical distribution of *F_IS_* is shown in filled purple. The asterisk at bottom left indicates the highest-weight model simulation (selfing rate = 100%, *N* = 100, *w_homozygote_* = lethal, mechanism = overdominance).

## 4. Discussion

We quantified patterns of genomic variation in *Corallorhiza bentleyi*, one of the rarest orchids in North America. Our analyses demonstrated distinctness between *C. bentleyi* and the Mexican *C. involuta*, supporting previous hypotheses that these are best treated as separate species (Barrett and Freudenstein, 2011). Analyses within *C. bentleyi* revealed weak patterns of population structure, characterized by inference of one ancestral population (*K* = 1). Surprisingly, we found evidence for excess heterozygosity across the genome (*H_O_* > *H_E_*) and associated negative values of inbreeding (*F_IS_*), which contradicts theoretical expectations for a rare, selfing species. Investigations of genomic diversity suggested elevated heterozygosity could not be explained by bioinformatic “noise,” small-*N* bias (Robertson Effect), or divergent, non-recombining chromosomal sequences (Meselson Effect). Demographic analyses strongly favored a model of post-LGM expansion without bottlenecks, and simulations under alternative selection scenarios supported overdominance vs. NFDS to explain excess heterozygosity and negative inbreeding.

### 4.1. *Corallorhiza bentleyi* individuals are predominantly cleistogamous or mixed chasmogamous-cleistogamous

Surveys revealed that cleistogamy is the predominant floral syndrome in *C. bentleyi*. While all localities had some chasmogamous flowers, typically in the form of mixed cleistogamy/chasmogamy within a single raceme, the higher prevalence of cleistogamy overall (coupled with seed set and ovary swelling early in floral development in virtually all flowers, regardless of chasmogamy) suggests that *C. bentleyi* is obligately self-pollinating (Fig. 1). Examination of herbarium records and iNaturalist observations of *C. involuta* yielded scant evidence for cleistogamy, although two iNaturalist observations appeared to be nearly cleistogamous. While difficult to rule out environmental or phenological effects from images, it was notable that the two most cleistogamous-appearing records occurred at the periphery of the known range of *C. involuta*. If indeed peripheral populations are cleistogamous or nearly cleistogamous, then it follows that *C. bentleyi*, isolated in eastern North America, would also be largely cleistogamous. Such patterns are observed in other plant species (e.g., Busch, 2005; Moeller, 2006; Cisternas-Fuentes et al., 2022), for example, in which selfing evolved as a response to a lack of suitable pollinators if niche spaces and geographic ranges of the plant and pollinator do not fully overlap.

### 4.2. *Corallorhiza involuta* and *C. bentleyi* are distinct, with weak population structure in *C. bentleyi*

Previous taxonomic treatments placed *C. involuta* as a variety within the *C. striata* complex (*C. striata* var. *involuta*) based on morphological similarities and molecular data (e.g., Freudenstein, 1997; Freudenstein and Senyo, 2008; Barrett and Freudenstein, 2009). However, these studies included few representatives of *C. involuta* and *C. bentleyi* or were based on few loci and lacked sufficient resolution. In the current study, *C. involuta* is differentiated from *C. bentleyi* (Figs. 3, 4), with *π* between the two species being greater than within each. While sampling of individuals was limited in *C. involuta*, the divergence between it and *C. bentleyi*, in addition to a fixed morphological character state (callus apex extension beyond widest point of the lip in *C. bentleyi* but not *C. involuta*; Barrett and Freudenstein, 2011), confirms species-level recognition following the Evolutionary Species Concept (Simpson, 1951; Wiley, 1978) as expounded by Freudenstein et al. (2017), though the exact ecological “role” of this morphological feature is unknown.

Given its life history as a rare, selfing, predominantly cleistogamous, mycoheterotrophic orchid with small population sizes, *C. bentleyi* was expected to display patterns reflecting those in apomictic “microspecies,” where localized asexuality creates unique evolutionary lineages (e.g., Grant, 1981; Barrett, 2003; Suetsugu et al., 2025). On the contrary, unsupervised analyses favored an optimal *K* = 1, revealing no population differentiation across sampled localities. Finer-scale analyses of population structure revealed subtle distinction among localities, particularly PM Monroe WV (Fig. 5), though we were unable to include genomic sampling of individuals from PC Pocahontas WV (Fig. 1). Our findings reflected those of the only previous study to include population-level sampling of *C. bentleyi*, both in overlapping genetic (dominant-marker ISSR) and morphological patterns (morphometrics of 14 floral characters; Fama et al., 2021). These patterns fit a history of post-glacial colonization from a refugial progenitor (e.g., Carnicero et al., 2022; Urquhart-Cronish et al., 2025). In fact, a somewhat similar, albeit more northern disjunction occurs within the *C. striata* complex, with populations of *C. striata* var. *vreelandii* patchily distributed throughout southwestern USA and Mexico, but disjunct to eastern Quebec and Newfoundland (Barrett and Freudenstein, 2009; 2011).

### 4.3. *Corallorhiza bentleyi* displays unexpected patterns of excess heterozygosity and negative inbreeding coefficients

The findings of excess heterozygosity and negative inbreeding run directly counter to our expectations for *C. bentleyi*. Cross-pollination by male ichneumonid wasps is known within the *C. striata* complex, to which *C. bentleyi* and *C. involuta* belong, but insect pollination (likely via sexual deception) has only been demonstrated in the larger, open-flowered *C. striata* var. *striata*, distributed in northern-montane USA and Canada (Freudenstein and Barrett, 2026). Given the obligately selfing reproductive mode of *C. bentleyi* (and *C. involuta* for that matter), it is unlikely that recent or sustained gene flow are plausible explanations for excess heterozygosity. Even if rare gene flow events do occur among individuals bearing a sufficient number chasmogamous flowers, they might only be expected to occur within sites, as the fragmented distribution of *C. bentleyi* could limit inter-population transfer of pollen or even successful dispersal of dust seeds (i.e., microscopic seeds lacking endosperm, characteristic of *Corallorhiza* and other mycoheterotrophic taxa; Rasmussen, 1995).

### 4.4. Excess heterozygosity and negative inbreeding are not explained by bioinformatic artifacts or small-*N* sampling

If bioinformatic “noise” caused excess heterozygosity (collapse of assembled paralogs), then we would expect to see a left-skewed distribution and peak of loci with “fixed” heterozygosity (near 100%), but we see the opposite (Fig. 6A). Assembly with a reference-free method (discoSNP) yielded similar findings of excess heterozygosity and negative *F_IS_*, arguing against pseudoreference artifacts as a cause for these findings (Fig. S2). Low levels of linkage disequilibrium (Table 4) argue against the Meselson Effect; this phenomenon is best studied in animals (e.g. Butlin, 2000; Schön, 2009; Danchin et al., 2011; Brandt et al., 2021), but has been demonstrated less frequently in plants (e.g., Pellino et al., 2013; Hojsgaard and Hörandl, 2015; Hernández-Velasco et al., 2025).

The negative relationship between *F_IS_* and minor allele frequency (*MAF*) is informative (Fig. 6C). When collapsed paralogs produce apparent heterozygosity, they are expected to affect loci randomly or bias toward intermediate frequencies (e.g., Gayral et al., 2013; McKinney et al., 2017; O’Leary et al., 2018). Even after removing loci with >85% *H_O_*, the negative relationship persisted, suggesting that excess heterozygosity is a genome-wide signal rather than a bioinformatic artifact (Fig. 6C). Our findings also suggest that excess heterozygosity is not merely the result of small sample sizes in some localities (Robertson Effect; Fig. 6D). Small sample sizes are expected to manifest as neutral sampling deviations (*H_O_* > *H_E_*) that generate negative *F_IS_*, but this bias may act in a non-linear fashion and is strongest at low *MAF*. Our finding of a linear decrease in *F_IS_* as *MAF* increases toward 0.5 is thus the opposite of the small-sample expectation (e.g., Robertson and Hill, 1984; Nei, 1987; Hartl and Clark, 2007).

### 4.5. Demographic modeling rejects bottlenecks despite widespread heterozygosity excess

The significant finding of excess heterozygosity relative to expectations under mutation-drift equilibrium could be attributed to a genetic bottleneck, but this test does not explicitly distinguish between the outcomes of drift vs. selection (Cornuet and Luikart, 1996). Demographic analyses with approximate Bayesian computation clearly favored a model of post-LGM divergence between *C. involuta* and *C. bentleyi* followed by range expansion, presumably from an ancestral glacial refugium (Table 6). Never was a model favored that included a genetic bottleneck, which may indicate small population sizes even ancestrally.

The most plausible explanation, based on our findings, is that selection maintains an excess of heterozygosity and thus apparent low inbreeding values. Specifically, selection against homozygous offspring might be expected to be significant, especially given the fact that fully mycoheterotrophic (i.e., non-photosynthetic) plants are known to have elevated mutation rates (e.g., Bromham et al., 2013; Lam et al., 2016; Barrett et al., 2018). This means that there may be a point at which selection pressure against homozygotes reaches an equilibrium with heterozygosity reduction due to obligate selfing (e.g., Workman and Jain, 1966; Brud, 2022). However, the efficacy of selection is diminished in selfing populations due to the strong effects of drift toward homozygosity (Glémin, 2007).

### 4.6. Simulations support overdominance over NFDS

Glémin (2021) discussed the outcomes of balancing selection in selfing species as the result of two potential selective processes that can maintain heterozygosity. First is overdominance, favoring heterozygotes and selecting against homozygotes. However, selective pressure is required to be intense to overcome the loss of heterozygosity due to selfing. A plausible way this could play out in *C. bentleyi* (and possibly *C. involuta*) is “purging” of low-fitness homozygous offspring early in development, so they never germinate or develop to maturity, which would represent the most extreme form of overdominance (homozygote lethality). This situation would be expected, however, to be associated with strong inbreeding depression in the sense that total reproductive output would be severely diminished (if homozygote fitness approaches zero; Husband and Schemske, 1996).

The second possible form of selection maintaining heterozygosity in selfing species is negative frequency dependent selection (NFDS). If selective pressures vary across space and/or time such that rare variants confer higher fitness, then heterozygosity could be maintained without overdominance, e.g., as has been demonstrated for pathogen resistance (e.g., Burdon and Thrall, 2009). Previous authors have hypothesized that if fungal host preference in *Corallorhiza* is under some degree of genetic control, then a signature of plant polymorphism associated with different fungal genotypes should be expected (e.g., Taylor et al., 2004; Fama et al., 2021). Alternatively, rare polymorphisms may be favored at different localities with different suitable fungal host genotypes, or over time as certain orchid-fungal genotype associations become predominant. In this sense, orchid genotypes could be “competing” for limited fungal resources, so a rare variant for even slightly different fungal genotype preferences could confer a selective advantage.

However, it will be challenging to tease apart genetic control and NFDS from broader plasticity of the orchids to associate with different fungal genotypes that happen to be more abundant in different sites or across time (e.g., forest succession or disturbance regimes). Experimental transfers of seed between localities could address this question but again, this risks depleting recruitment at any one source site, given how few flowering individuals are present at a site in a given year. Regardless, our simulations do not provide evidence for the role of NFDS in excess heterozygosity (Fig. 7; Table S1). Seed viability assays and germination trials (e.g., Pritchard, 1985; Wood and Pritchard, 2004), coupled with genotyping and assessment of heterozygosity levels, would be particularly informative (but challenging due to microscopic seed size) in determining the relationship between heterozygosity and fitness. In other words, it would help determine the fraction of viable seeds from a single capsule, each of which can produce thousands of minute “dust seeds,” and how this relates to heterozygosity.

### 4.7. Conservation implications

Our inference that overdominance and “lethal” homozygosity provide the most likely among alternative explanations for excess heterozygosity in *C. bentleyi* has implications for reproductive output and extinction risk. If homozygosity truly is lethal (*w_homozygote_* ∼ 0), then reproductive output should be cut in half relative to neutral expectations. It is well demonstrated that mycoheterotrophs, and particularly many orchid taxa, are extremely sensitive to forest disturbance (e.g., Merckx, 2013; Crain et al., 2025; Watkinson, 2025). This means that population recovery from local, regional, or global impacts may be significantly hindered, putting plants like *C. bentleyi* in “double jeopardy,” with both intrinsic and extrinsic environmental threats. This fragility is borne out by our field observations; seven annual surveys (2016-2026) of the largest historically known population in Giles County, Virginia, yielded zero racemes (likely associated with selective logging prior to 2016), and is possibly extirpated.

The primary conservation priority is to minimize habitat degradation. Most known *C. bentleyi* sites occur near roads or trails at the transition of disturbed and maturing second-growth forests; the holotype locality is on an abandoned logging railroad bed converted to a recreational trail (Freudenstein, 1999; Bentley, 2001; 2004; Fama et al., 2021). This means nearly all known sites are at risk of anthropogenic disturbance such as logging, pollution, pesticides, human traffic, and invasive species. Stiltgrass [*Microstegium vimineum* (Trin.) A.Camus], introduced from Asia, has invaded several *C. bentleyi* sites, where avoidance of this high-abundance grass by herbivores intensifies grazing pressure on native species (e.g., Knight et al., 2009). Further, nearly all sites lie within mixed-use National Forest but not within designated Wilderness Areas, so the threat of logging presents a risk, as these mycoheterotrophic orchids are especially sensitive to forest structural disturbance. In this sense, *C. bentleyi* is also indirectly at risk from forest pests that affect the survival of surrounding trees, the sources of photosynthate via ectomycorrhizal intermediaries (e.g., Cheaib and Killiny; 2025).

Addressing these conservation challenges is further hindered by scarcity of basic ecological data for *C. bentleyi*. Because the species is known from only approximately a dozen sites across a narrow 200-km Appalachian range, conventional environmental niche modeling approaches lack sufficient occurrence points to generate reliable habitat suitability maps (van Proosdij et al., 2015). Instead, identifying suitable microhabitats or potential translocation sites requires targeted surveys with soil metabarcoding to confirm the presence of suitable fungal genotypes within the family Thelephoraceae; more specifically, within the few species of the genus *Thelephora* Ehrh. ex Willd. (formerly *Tomentella* Pers. ex Pat.; Barrett et al., 2010; Fama et al., 2021; Barrett et al., 2022) upon which *C. bentleyi* depends. However, conducting *ex situ* propagation or field seed-sowing experiments with *C. bentleyi* poses a demographic risk. Harvesting seed capsules from wild populations can deplete local seedling recruitment in years when few racemes emerge, exacerbated by reduced potential overall seed viability stemming from lethal homozygosity. To mitigate this risk, more abundant and co-occurring congeners, such as *C. maculata* var. *maculata* (Raf.) Raf. (eastern spotted coralroot) and *C. odontorhiza* (Willd.) Poir. (autumn coralroot), could serve as experimental proxies. Optimizing protocols for symbiotic seed germination, fungal host-matching, and microbial detection in these more abundant and widespread congeners may establish a benchmark before applying management interventions to *C. bentleyi*.

## 5. Conclusion

While our findings of widespread excess heterozygosity were unexpected, they raise a broader question: how many rare taxa with similar reproductive modes are able to maintain comparable genomic architectures? Among orchids, an estimated 88% of species are self-compatible.

Globally, approximately 15-20% of the family relies on autonomous self-pollination (Ackerman et al., 2023). Extrapolating across the estimated 30,000 species of Orchidaceae, over 4,600 orchid species may autonomously self-fertilize (representing nearly 1.3% of all angiosperms), suggesting that, while in the minority, selfing is not a mere evolutionary dead-end but a recurrent and pervasive strategy (e.g., *Gastrodia*; Suetsugu et al., 2025). Because many selfing orchids are rare or threatened, the family potentially serves as a model system for addressing this apparent reproductive-genomic paradox. Although our simulations favored overdominance to explain excess heterozygosity in *C. bentleyi*, experimental validation of fitness across life stages (fecundity, seed viability, protocorm development, and adult survival) remains key to confirming the links between modes of selection and long-term persistence. Habitat fragmentation and insect-pollinator population declines may force rare plant species toward self-fertilization; thus, understanding genomic “strategies” to mitigate homozygosity is of importance to biodiversity conservation. Investigation of how *C. bentleyi* and other rare, selfing species might leverage heterozygote advantage to buffer against the most severe impacts of inbreeding depression has potential to improve our understanding of genetic rescue, revealing that genome-wide overdominance may be an intrinsic mechanism to shield selfing species from extinction.

## 6. Acknowledgements

Funding was provided by the US National Science Foundation award DEB CAREER: 2044259 to CFB; the West Virginia University Department of Biology and Eberly College; and by the West Virginia Research Challenge Fund through a grant from the Division of Science and Research, HEPC and in part by (i) the WVU Provost’s Office, (ii) the Eberly College of Arts and Sciences, (iii) the Honors College and (iv) the Department of Biology to NMF. We thank Stanley Bentley, Lidia Cabrera, Charles Garratt, Nancy Van Alstine, and the USDA Forest Service for assistance in the field. All sampling was conducted with permission from and according to the guidelines stipulated by the US Department of Agriculture Forest Service and the Secretaría de Medio Ambiente y Recursos Naturales (SEMARNAT) Mexico. Material was transferred from Mexico to the United States via a CITES Scientific Exchange Certificate between Herbario Nacional de México (herbarium code MEXU) and the Ohio State University Herbarium (code OS). We thank Ryan Percifield for sequencing support and discussion, and the WVU Genomics Core Facility for support provided to help make this publication possible and CTSI Grant #U54 GM104942 which in turn provides financial support to the WVU Core Facility.

## 8. Data Accessibility and Benefit-Sharing

Raw sequence reads are deposited in the US National Center for Biotechnology Information Sequence Read Archive (SRA) under BioProject PRJNA771539. Genomic SNP datasets, metadata, and code are available on Zenodo (https://doi.org/10.5281/zenodo.23220344). All code used in this work is also available via GitHub (https://github.com/barrettlab/C_bentleyi_2026).

Google Gemini Flash v.3.8 (Google, 2026) was used to troubleshoot UNIX and R code, and to help build the simulation code for this study. *Benefit sharing*: A research collaboration was developed between scientists from Mexico and the USA. All collaborators are included as co-authors, and the results of research have been shared openly and freely with the provider communities and the broader scientific community (see above). The research addresses a priority concern, in this case the conservation of the organisms being studied. More broadly, our group is committed to international scientific partnerships, as well as institutional capacity building.

## 9. Author Contributions

CFB, NMF, SDS and BTS conceived of the study. CFB, JVF, and GS conducted fieldwork. CFB, BTS, SDS and NMF conducted lab work and analyzed data. CFB, BTS, and SDS developed data analysis code. CFB led writing and all coauthors contributed to drafts of the manuscript.

---

**Table S1.** Simulation-based tests of alternative selection scenarios maintaining excess heterozygosity and negative inbreeding coefficients (*F_IS_*). ‘*N*’ = population size (at a single locality), ‘Selfing’ = selfing rate, ‘Selection’ = mode of selection (OD = overdominance; NFDS = negative frequency dependent selection); ‘Intensity’ = a summary of the relative fitnesses of the three genotypes at each locus; ‘W-distance’ = Wasserstein distance between the each simulation-based and the empirical *F_IS_* distribution; ‘Weight’ = the relative model weight.

| $N$ | Selfing | Selection | Intensity | W-distance | Weight |
| --- | --- | --- | --- | --- | --- |
| 100 | 100% | OD | lethal | 0.1118592 | 0.193717 |
| 100 | 90% | OD | lethal | 0.1188989 | 0.188208 |
| 10 | 90% | OD | lethal | 0.1400001 | 0.170806 |
| 10 | 100% | OD | lethal | 0.1650596 | 0.149121 |
| 10 | 90% | OD | strong | 0.2172917 | 0.104592 |
| 100 | 90% | OD | strong | 0.2360588 | 0.089924 |
| 100 | 100% | OD | strong | 0.2611567 | 0.072048 |
| 10 | 100% | OD | strong | 0.3395588 | 0.031215 |
| 10 | 90% | OD | moderate | 0.6340444 | 0.000192 |
| 100 | 90% | OD | moderate | 0.6561319 | 0.000116 |
| 10 | 90% | OD | neutral | 0.6892754 | 5.24E-05 |
| 10 | 90% | NFDS | strong | 0.7991004 | 2.87E-06 |
| 100 | 90% | NFDS | strong | 0.8101258 | 2.1E-06 |
| 10 | 90% | NFDS | weak | 0.8347374 | 1.02E-06 |
| 100 | 90% | OD | weak | 0.8420975 | 8.21E-07 |
| 10 | 90% | OD | weak | 0.8451775 | 7.48E-07 |
| 100 | 90% | NFDS | weak | 0.8586302 | 4.98E-07 |
| 100 | 90% | OD | neutral | 0.8682218 | 3.71E-07 |
| 100 | 100% | OD | moderate | 1.0531936 | 6.73E-10 |
| 10 | 100% | NFDS | strong | 1.0566358 | 5.92E-10 |
| 10 | 100% | NFDS | weak | 1.0566358 | 5.92E-10 |
| 10 | 100% | OD | moderate | 1.0566358 | 5.92E-10 |
| 10 | 100% | OD | neutral | 1.0566358 | 5.92E-10 |
| 10 | 100% | OD | weak | 1.0566358 | 5.92E-10 |
| 100 | 100% | NFDS | strong | 1.0566358 | 5.92E-10 |
| 100 | 100% | NFDS | weak | 1.0566358 | 5.92E-10 |
| 100 | 100% | OD | neutral | 1.0566358 | 5.92E-10 |
| 100 | 100% | OD | weak | 1.0566358 | 5.92E-10 |

**Figure S1.**
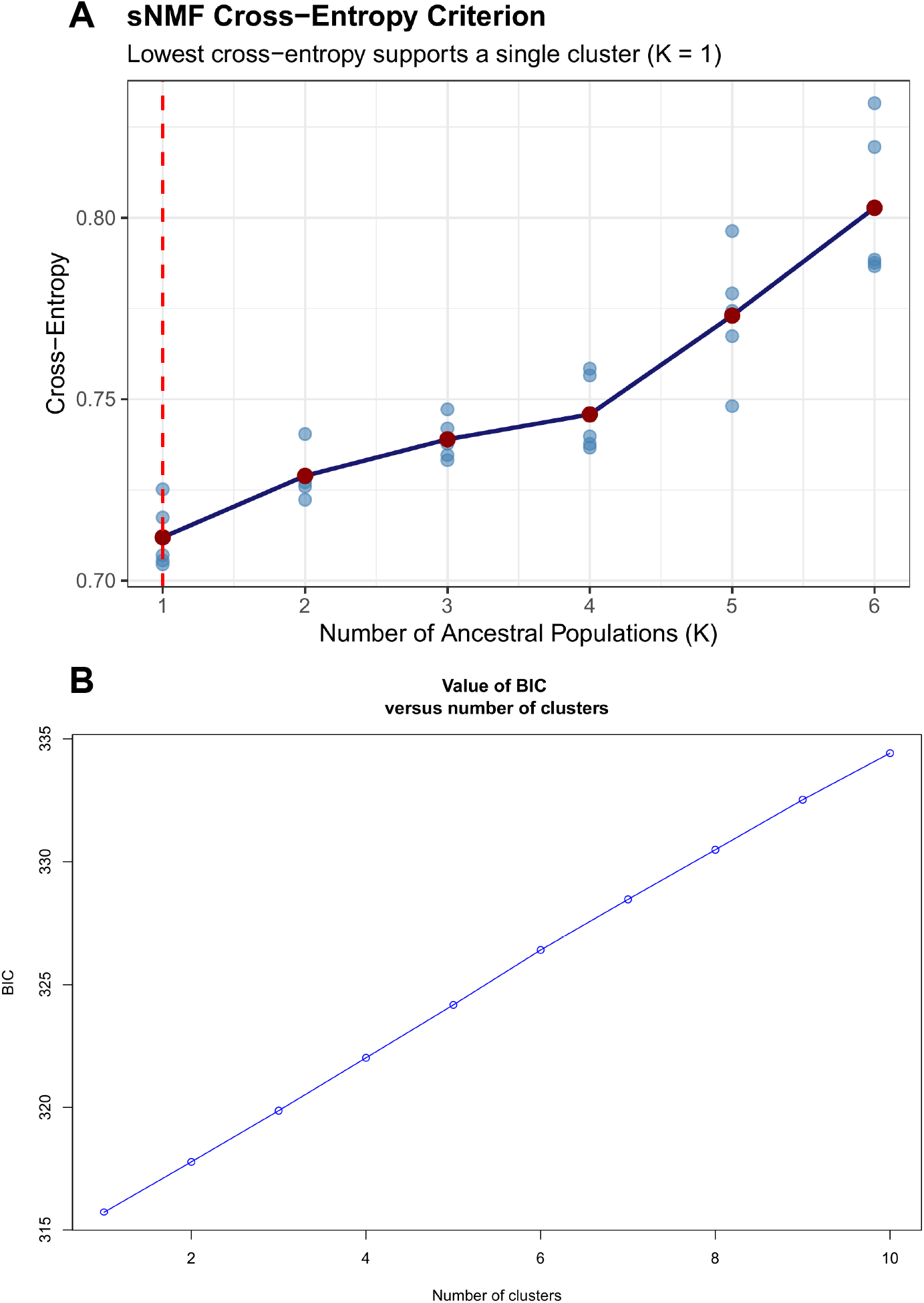
Unsupervised population structure assessment of the optimal number of ancestral population clusters (*K*). **A**. Cross-entropy values for K = 1-6 based on unsupervised sNMF in LEA. **B**. Bayesian Information Criterion (BIC) for K = 1-10 in unsupervised DAPC.

**Figure S2.**
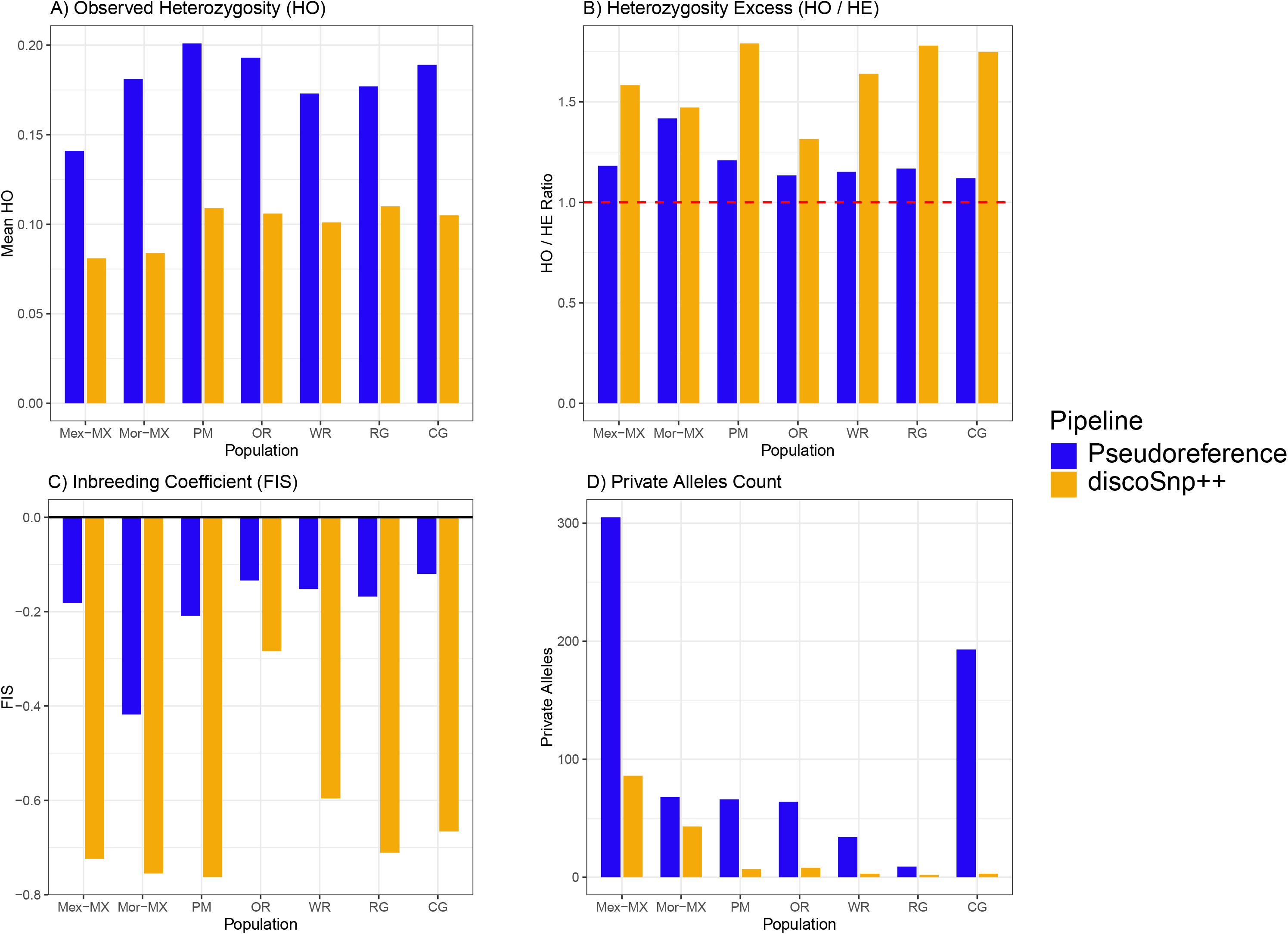
Comparison of diversity metrics between the original pseudoreference assembly and a reference-free assembly with discoSNP++.

